# Regulation of heterosis-associated gene expression complementation in maize hybrids

**DOI:** 10.1101/2024.10.30.620956

**Authors:** Marion Pitz, Jutta Baldauf, Hans-Peter Piepho, Peng Yu, Heiko Schoof, Annaliese S. Mason, Guoliang Li, Frank Hochholdinger

## Abstract

The dominance model of heterosis explains the superiority of F_1_-hybrids via the complementation of unfavorable by beneficial alleles in many genes. Consistent with this model, genes active in only one parent and the hybrid display single-parent expression (SPE) complementation. Here we demonstrated, that SPE can explain up to 29% of heterotic variance in maize (*Zea mays* L.). Moreover, expression quantitative trait loci (eQTL) revealed, that the genotype of the active parent determines the regulation mode of SPE patterns in *cis* or *trans*, which might explain how phylogenetic distance translates to transcriptomic diversity and heterosis in hybrids. Furthermore, we showed that most eQTL are located in heterozygous regions of the genome and that a *trans* eQTL controls the activity of a SPE gene which regulates lateral root density in hybrids. We anticipate that our study will stimulate further research elucidating the regulation and molecular mechanisms underlying heterosis.

## Introduction

Heterozygous F_1_ hybrids are more vigorous and show higher fitness and biomass production compared to their homozygous parental inbred lines^1^, a phenomenon called heterosis^2^. The introduction of hybrids in plant breeding was one of the landmark innovations of agriculture. Among crops, outcrossing species such as maize display the highest degree of heterosis^3^.

Heterosis is typically measured for above-ground traits related to yield and biomass^4^, but root traits, which have been largely neglected in the past, are also important for future crop improvement^5^. In maize roots, heterosis manifests as early as 5 to 7 days after germination for traits such as primary root length or lateral root density^6^.

One classical genetic mechanism to explain heterosis is the dominance model, which assumes that many slightly deleterious alleles are complemented in the hybrid by dominant or at least stronger alleles^7^. On the gene expression level, single parent expression (SPE) complementation, where the hybrid expresses a gene that is only active in one of its parents is consistent with the dominance model^8,9^. SPE complementation results in hundreds of additionally active genes in hybrids. It probably plays a role in translating parental diversity into phenotypic heterosis and has been linked to hybrid vigor^10^. It was further established that SPE genes are significantly enriched among evolutionarily younger, non-syntenic genes and might function in the adaptation of hybrids to different environments^10–12^. Additionally, variation in transcriptional regulation of *cis*- and *trans*-acting factors has been highlighted in relation to hybrid performance^13^. *Trans*-regulated gene expression in hybrids was associated with paternal alleles in maize^14^. Moreover, an association between SPE and *cis*-regulation was suggested^12^.

Intermated recombinant inbred lines (I-RILs) are the result of crossing two distinct inbred lines, followed by several generations of intercrossing and subsequent self-pollination of their progeny^15^. In maize, the intermated B73×Mo17 recombinant inbred line (IBM-RIL) syn. 4 population, a set of ~300 highly homozygous intercrossed RILs (4 generations of intercrossing), shows a high diversity of phenotypes and of genomic regions contributed by the two parental genotypes^16^. By backcrossing RILs to their original parents, backcross populations can be generated which show varying degrees of heterozygosity and heterosis. Genetically and phenotypically diverse RILs and backcross populations are an important resource for QTL mapping, as well as for candidate gene identification and heterosis studies^17–20^.

We used the IBM-RIL population and two backcross populations to study how varying heterozygosity and the regulation of SPE complementation influences the manifestation of heterosis in seedling root development. We demonstrated, that SPE genes are predominantly regulated from heterozygous regions of the genome and that depending on the genetic constitution of the active parent, *cis*- or *trans*-regulation of SPE is prevalent. We hypothesize based on our findings, that differences in regulation of the parental lines of a hybrid contribute to heterosis.

## Material and Methods

### Plant genetic resources

The IBM-RIL syn. 4 population^16^ represents a collection of highly diversified genotypes with respect to the genomic regions contributed by their two parental inbred lines B73 and Mo17 (Figure 1A). To study the phenotypic and transcriptomic plasticity of maize F_1_ hybrids relative to their parents, a random subset of 112 IBM-RILs, was backcrossed to their original parental inbred lines B73 and Mo17. In all crosses, B73 and Mo17 were selected as the female parent to secure a homogenous phenotype of all plants on which the pollinated ears will develop and thus similar seed quality. Each of the two F_1_-backcross hybrids of a specific IBM-RIL (hereafter named B73×IBM-RIL and Mo17×IBM-RIL backcross hybrids) show contrasting homozygous and heterozygous genomic regions and genes (Figure 1A). For example, if a region between two recombination breakpoints is homozygous in a B73×IBM-RIL backcross, it is heterozygous in the corresponding Mo17×IBM-RIL backcross hybrid and vice versa.

**Figure 1:**
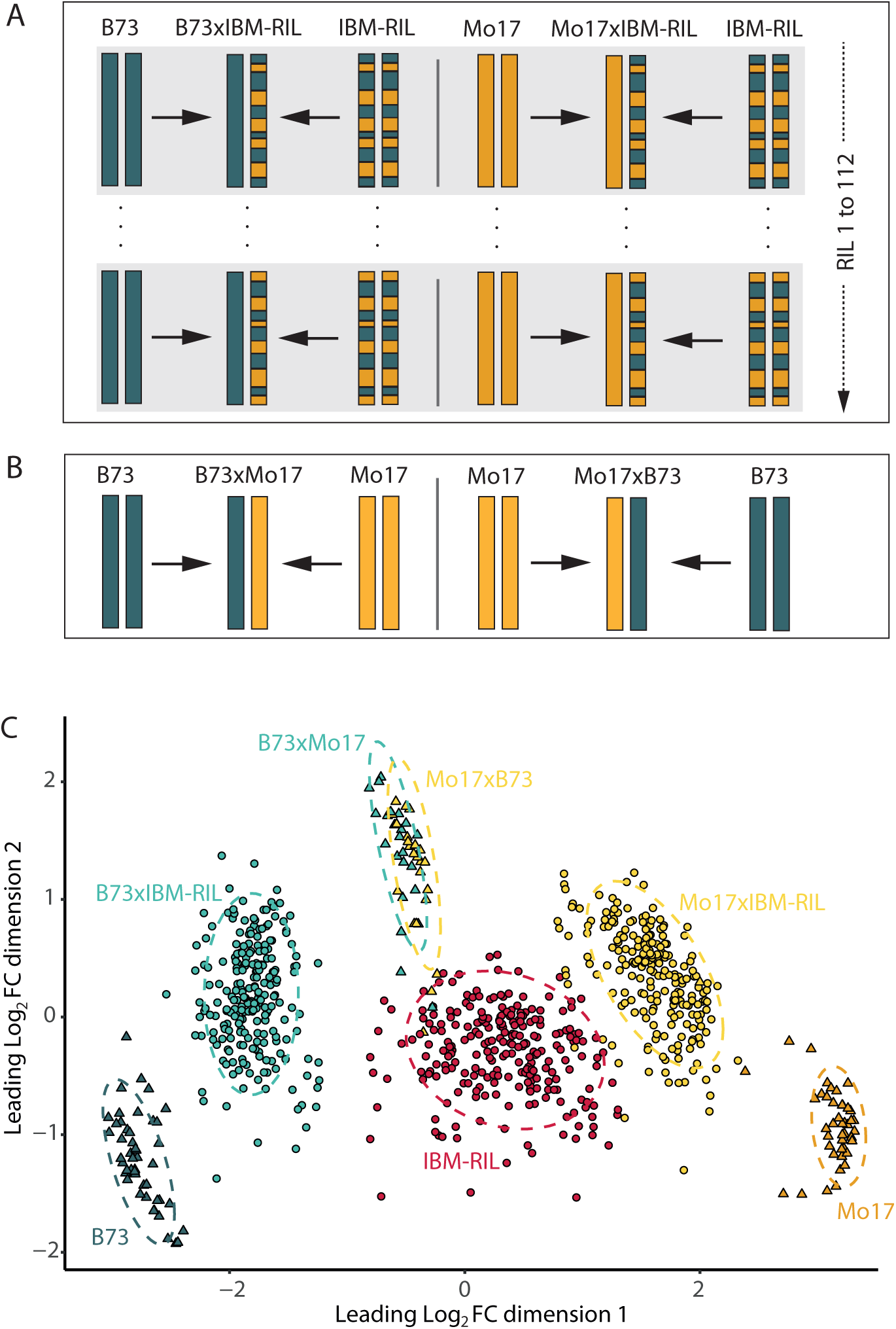
Plant genetic material and sample information. **A** Schematic depiction of the 112 IBM-RILs and their B73- and Mo17-backcross hybrids. **B** Schematic depiction of the reference genotypes. **C** Multidimensional Scaling Plot of 834 high-quality RNA-seq samples. Each genotype group is highlighted by a different colour. The reference genotypes B73, Mo17, B73xMo17 and Mo17xB73 are depicted as triangles, IBM-RILs and their backcrosses as circles. The leading Log_2_ FC between two samples can be interpreted as expression difference between those two samples.

### Experimental design

We studied the phenotypic and transcriptomic plasticity of the IBM-RIL backcross hybrids relative to their parental inbred lines. A selection of 112 IBM-RILs were used as paternal inbred lines, corresponding to 112 B73×IBM-RIL and 112 Mo17×IBM-RIL backcross hybrids (Figure 1A). To optimally fit the experimental design and increase the precision of subsequent pairwise comparisons with the common parental inbred lines and reference hybrids both common maternal inbred lines B73 and Mo17 and the two reference hybrids B73×Mo17 or Mo17×B73, respectively, were included (Figure 1B). For each sample, 25 kernels of the same genotype (parent or hybrid) were surface sterilized in 10% H_2_O_2_ for 20 min, rinsed with distilled water and afterwards pre-germinated in filter paper rolls with five kernels each in a climate chamber with a 16 h light (26 °C), 8 h dark (21 °C) cycle in distilled water^11^. After three days, eight seedlings per genotype with approximately the same length of primary root and, if already present, shoot length were selected and transferred into a row of an aeroponic growth system for four additional days. Each aeroponic growth system (“Elite Klone Machine 96”, TurboKlone, USA) was composed of 12 rows each with eight planting sites. Thus, we could fit 12 different genotypes into one aeroponic growth system and eight systems at the same time into our climate chamber (16 h light, 26 °C; 8 h dark, 21 °C) (Figure S1). We analyzed three independent biological replicates, each comprising all IBM-RIL inbred lines and hybrids. Due to the large number of samples and space limitations, the different genotypes of each biological replicate were grown in four batches distributed across four weeks, also called alpha-design with incomplete blocks^21^. Within each batch, the eight aeroponic growth systems were randomly assigned to eight positions in the climate chamber. Three successive rows of an aeroponic growth system represent one triplet. To each triplet an IBM-RIL and its corresponding B73 and Mo17 backcross hybrids, or both common maternal inbred lines B73 and Mo17 and one of the two reference hybrids B73×Mo17 or Mo17×B73, respectively were assigned. The randomization process was conducted at the replicate level for the triplets, whereas it was ensured, that in each batch two reference triplets each of B73, Mo17, B73xMo17 and B73, Mo17, Mo17xB73 were distributed. Thus, in each batch we surveyed 30 IBM-RIL triplets and both reference hybrids and the common inbred lines B73 and Mo17 in two additional triplets (Figure S1). So that in total, 3 samples of each IBM-RIL and each backcross hybrid, 48 samples (biological replicates) of the maternal inbred lines B73 and Mo17, and 24 samples of the reciprocal hybrids B73×Mo17 and Mo17×B73 as reference hybrids were analyses. In other words, each independent biological replicate contained 384 samples (in total: 384 x 3 replicates = 1,152 samples). The number of individual samples per replicate was designed to also fit one sequencing run on the NovaSeq 6000 S4 flow cell machine (Illumina, San Diego, USA), described later.

### Root phenotyping and sampling for RNA-seq

Seven days after germination, all seedlings per sample (maximum eight seedlings) were removed from the aeroponic growth system and the seedling root system was scanned using an Epson Expression 12000XL scanner (Epson, Meerbusch, Germany) with up to four plants per image. The resulting images were cropped to create single plant images that only showed the root system and the maize kernel. We used the RootPainter software client (version 0.1.0) and server component (version 0.2.7) to train a convolutional neural network to recognize and segment roots in images^22^. We then analyzed the segmented images in a batch using RhizoVision Explorer (version 2.0.3)^23^. After inspection and cleanup, we determined the total root length, total root volume and number of root tips for each plant for subsequent analysis (Details in Supplement Material SM1).

After imaging the seedlings, the primary root was separated from the kernel to collect (i) the proximal first centimeter with emerged lateral roots in 80% Ethanol to count the number of lateral roots per cm as density and (ii) the distal region of the primary root, composed of the root tip and the meristematic zone followed by the elongation zone, in liquid nitrogen for subsequent RNA extraction.

### Analysis of phenotypic data

To evaluate the phenotypic data of each genotype a linear mixed model (baseline model 1) with a fixed effect for block (three replicates as levels) and genotype (263 levels) was fitted. According to the layout of our experimental design (Figure S1) we included random effects for row, triplet, system and batch effect in the model. The residual error assesses the within-row variance among plants.

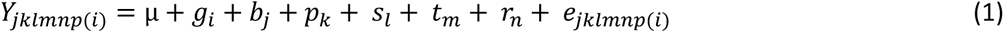

*Y_jklmnp_*_(*i*)_ represents the mean phenotypic value of a specific trait of interest of the respective genotype *i*, *µ* represents the intercept, *g_i_* represents the fixed effect for genotype *i*, *b_j_* represents the fixed effect for block *j*, *p_k_* represents the random effect for batch *k* nested within block *j*, *s_l_* represents the random effect for system *l* nested within batch *k* and block *j*, *t_m_* represents the random effect for triplet *m* nested within system *l*, batch *k* and block *j*, *r_n_* represents the random effect for row *n* nested within triplet *m*, system *l*, batch *k* and block *j*, and *e_jklmnp_*_(*i*)_ represents the random error effect for plant *p* of genotype *i* in block *j*, batch *k*, system *l*, triplet *m* and row *n*. To fulfil the assumptions of linear models, the phenotypic values for the traits “total root length”, “total root volume” and “total number of root tips” had to be square root-transformed. An offset of 0.5 was added to each phenotypic value before transformation. The resulting modelled means were transformed back to their original scale for visual inspection (Figure S3). Modelled means on the transformed scale (and original scale in case of lateral root density) were used for TWAS analysis (described below).

### RNA-sequencing and preparation of alignments

For subsequent RNA extraction and sequencing, a maximum of eight primary roots of each genotype grown in the same row of an aeroponic system were pooled. These root samples were manually ground in liquid nitrogen and total RNA was isolated with the RNeasy Plant Mini Kit (QIAGEN, Venlo, the Netherlands). RNA quality was assessed with a Bioanalyzer (RNA ScreenTape + TapeStation Analysis Software 3.2, Agilent Technologies, Santa Clara, CA, USA) by the Next Generation Sequencing (NGS) Core Facility in Bonn, Germany (https://btc.uni-bonn.de/ngs/), which subsequently constructed cDNA libraries for RNA-seq according to the TruSeq stranded mRNA library preparation protocol (Illumina, San Diego, USA). Sequencing was performed on a NovaSeq 6000 S4 flow cell machine (Illumina, San Diego, USA), generating 100-bp paired-end reads. This allowed for processing all 384 samples of a single replicate in one flow-cell and each batch of 96 samples on one lane. The obtained reads are reversely stranded. The raw reads were trimmed and filtered using Trimmomatic (version 0.39) in paired-end mode with the following settings: ILLUMINACLIP:adapters/TruSeq3-PE-2.fa:2:30:10:8:True, LEADING:3, TRAILING:3, MAXINFO:30:0.8, and MINLEN:40. With this step, remaining adapter sequences were removed, low quality bases from the start and end of the reads were cropped and adaptive quality trimming was performed. After these quality control, reads with a minimum length of 40 bases were retained and resulting single-end reads were excluded^24^. The maize reference genome B73 version 5 (B73v5, ftp.ensemblgenomes.org/pub/plants/release-52/fasta/zea_mays/dna/Zea_mays.Zm-B73-REFERENCE-NAM-5.0.dna.toplevel.fa.gz) was indexed with exon information from the corresponding annotation file (http://ftp.ensemblgenomes.org/pub/plants/release-52/gff3/zea_mays/Zea_mays.Zm-B73-REFERENCE-NAM-5.0.52.gff3.gz). The trimmed reads were aligned to the indexed reference genome using Hisat2 (version 2.2.1)^25^ with the appropriate input file settings and intron lengths: -q –phred 33 –rna-strandedness RF --min-intronlen 20 –max-intronlen 60000. The data was then saved in BAM format using the samtools view command from htslib (version 1.14)^26^. Picard tools (version 2.27.1; http://broadinstitute.github.io/picard/) was used to remove duplicates using MarkDuplicates.

Reads aligned to exons of genes were counted using htseq-count (version 2.0.1), with specifications to only count uniquely mapped reads^27^. Samples with less than 5 million counted reads were excluded.

### Preparation of alignments for SNP calling

For SNP calling between the genotypes of this study and the B73 reference genome, the read alignments were processed using the HaplotypeCaller of GATK (version 4.2.6.1) with respect to GATK’s best practices for RNA-seq data (https://gatk.broadinstitute.org/hc/en-us/articles/360035531192-RNAseq-short-variant-discovery-SNPs-Indels-, checked on 12/14/2023). First, Picard’s AddOrReplaceReadGroups (version 2.27.1) was used to add readgroup information to the alignments of all samples. The replicate number was set as RGLB, the RGPL field was set to ‘ILLUMINA’, the RGPU field was set to ‘unknown’, and the RGSM field was filled with the sample name. The samtools view (version 1.14) command was used to filter for uniquely mapped reads by only including reads with mapping quality of 60 or higher and to format and index the alignments. Second, GATK’s SplitNCigarReads was used to split alignments at positions with N in the CIGAR field, such as intron-spanning alignments^28^.

### SNP calling between B73 and Mo17 samples and B73v5 reference for sample evaluation

In brief, the GATK HaplotypeCaller was used to identify variants between the Mo17 samples and the B73v5 reference genome. The frequency of the B73 and Mo17 alleles at each SNP locus was previously identified in a similar manner^29,30^. The ratio of homozygous loci was calculated and samples with less than 95% homozygosity across expectedly homozygous loci were excluded (details in Supplementary Material SM2). A total of 175 RNA samples were excluded because they did not meet the criteria of being homozygous in ≥95% of the supposedly homozygous loci. Additionally, 10 samples were excluded beforehand due to their library size being <5 million read counts. Moreover, 17 samples were excluded because only one out of three replicates was left for the respective genotype. Since downstream analyses include the comparisons between both parents and their resulting hybrids, we had to further exclude 90 hybrid RNA-seq samples because all corresponding paternal IBM-RIL RNA-seq samples were excluded. In addition, 8 IBM-RIL RNA-seq samples were excluded because the corresponding hybrid samples were missing. Finally, 852 RNA-samples remained for subsequent SNP calling as described below (Data file S1).

### SNP calling between all high-confidence samples and the B73v5 reference

In the second SNP calling, SNPs between each sample and the B75v5 reference were called. We included the previously identified variants between our Mo17 samples and the B73v5 reference, as well as variants from our B73 samples vs. the B73v5 reference. They were filtered based on several criteria. The mapping quality (MQ), variant site quality (QUAL), Fisher strand (FS) and allele depth (AD) of SNP alleles were used with different thresholds for InDels and SNPs with respect to GATKs’ guide on hard-filtering short variants (https://gatk.broadinstitute.org/hc/en-us/articles/360035890471-Hard-filtering-germline-short-variants, checked on 12/18/2023). For the SNPs the filters QD >2, SOR <3, MQ >40, QUAL >30, FS <60 and FORMAT/AD[0:1] >5 were applied. For the indels, the filters QD >5, QUAL >30, FS <200 and FORMAT/AD[0:1] >5 were applied. The base qualities of each sample were then recalibrated using the filtered SNPs and indels as known-sites with GATK (version 4.2.6.1). BaseRecalibrator was run to generate recalibration tables, which were then applied to the aligned reads with ApplyBQSR (https://gatk.broadinstitute.org/hc/en-us/articles/360035531192-RNAseq-short-variant-discovery-SNPs-Indels-, checked on 12/14/2023). The HaplotypeCaller was run with the recalibrated samples in BP_RESOLUTION mode and reported the SNP sites at each individual position. The resulting variant files were then filtered for positions with a coverage (DP) of ≥1, to eliminate loci without any information. The variant files from Mo17 and B73 samples were combined by GenomicsDBImport. The samples of a triplet (IBM-RIL samples plus corresponding B73×IBM-RIL and Mo17×IBM-RIL hybrids) were combined with the Mo17 and B73 samples, resulting in one database per triplet. Genotyping of all samples within each database was performed using the GenotypeGVCFs function. Since the Mo17 and B73 samples are present in each database, we ensured that genotyping was performed on the loci differentiating B73 and Mo17 in each database^28^. The genotyping data was then filtered for SNPs with QD >2, SOR <3, MQ >40, QUAL >30 and FS <60 using bcftools (version 1.17). A list of high confidence SNPs was created in R using the results from the HaplotypeCaller of the Mo17 and B73 samples (Data file S2). For these loci, it was established that the genotyping results of the HaplotypeCaller (B73v5 reference allele vs. non-reference allele) correspond to the B73 and Mo17 alleles of the germplasm of this study (reference = B73, non-reference = Mo17): The HaplotypeCaller reports for each SNP locus of each sample the most likely genotype, which we term genotype-call in the following and the corresponding genotype quality (GQ), a measure for the confidence of the genotype-calls. Only genotype-calls of bi-allelic loci with a GQ of ≥10 were considered. The loci were filtered to include only those where ≥90% but a minimum of three remaining genotype-calls in Mo17 samples are homozygous for the non-reference allele, and 90% but a minimum of three remaining genotype-calls in B73 samples are homozygous for the reference allele. Next to the high confidence B73 vs Mo17 SNPs, we identified SNPs which did not belong to B73 or Mo17 as IBM-RIL specific (homozygous or heterozygous and regardless of GQ). Loci genotyped with a GQ of <10 were filtered. Only loci that were either in the high confidence SNP list of B73 vs. Mo17 alleles or which were IBM-RIL-specific (for masking putative IBM-RIL specific regions) are considered further.

### Classification of IBM-RIL genomic regions

The filtered SNP data were used to classify each IBM-RIL genome into B73 or Mo17 regions and to mask regions which were not B73 or Mo17. A distance-function was used to calculate the distance between the IBM-RIL specific loci. Loci with a distance of <2.5 Mbp were grouped together as a block. Blocks containing a minimum of 10 IBM-RIL-specific loci, with ≥5 of those being homozygous, were identified and masked as IBM-RIL-specific third origin regions. The start and end positions of these regions were recorded and loci within those regions were dropped. Next, a sliding window approach was used to eliminate singular loci that did not match their surrounding loci. A window of 15 loci was used, and ≥11 had to be homozygous for the Mo17 allele for the window to be considered a Mo17 window. For a B73 window, ≥12 out of 15 loci must have homozygous B73 alleles. The values for the windows were obtained by computing the minimum number of matching loci in a 15-loci window across B73 and Mo17 samples. Otherwise, the window was considered ambiguous^31^. Loci within an ambiguous window were dropped, as well as loci which were classified differently from their window. The previously mentioned distance-function was utilised to calculate the distance between the remaining loci. Loci that carried the same allele and which were less than 0.5 Mbp apart were grouped together as a block, and all blocks were retained. The start and end positions of these blocks were recorded as the Mo17 and B73 regions within each IBM-RIL (Data file S3). Two IBM-RILs which had more than 50% of their genomes consisting of IBM-RIL specific regions from a third parental origin were excluded along with their hybrids (Data file S1), leaving 834 samples for final analyses. The data set of each triplet reported by the HaplotypeCaller was filtered to only include loci within the B73 or Mo17 regions of the IBM-RILs and within exons of protein-coding genes. We checked for all protein-coding genes whether they were located in a B73 or Mo17 region or masked as neither a B73 or Mo17 region, or whether they were located in a genomic region without SNP information. This verification was performed for each IBM-RIL separately. Centromere locations of the 10 chromosomes were taken from the genome assembly of MaizeGDB by selecting the “Knobs, centromeres and telomeres” information https://jbrowse.maizegdb.org^32^. The proportion of heterozygous to homozygous regions was calculated for each backcross hybrid by dividing the total lengths of classified heterozygous regions (B73 regions of the IBM-RIL for the Mo17×IBM-RIL and Mo17 regions of the IBM-RIL for B73×IBM-RIL) by the total lengths of all classified regions (not considering IBM-RIL specific masked regions and regions without SNP information).

### Multidimensional Scaling (MDS)

To evaluate the quality and the structure of the RNA-seq samples in this study, a multidimensional scalig (MDS) plot was used. The active genes of the 834 filtered samples were compared by the plotMDS() Bioconductor package limma (version 3.50.3)^33^ in R.

### Analysis of expression complementation

The activity/expression status of each gene was determined as previously described^10^ based on thresholding normalized read counts. In short, fitting a generalized additive model (R package mgcv)^34^ using guanine-cytosine (GC) content and log-transformed gene length as explanatory variables to account for artifactual read count differences across genes^35^ resulted in a predictive count for each gene. The inverse of the predictive count was used as a multiplicative gene-specific normalization factor. In addition, sample-wise scale factors using the trimmed mean of M-values (TMM) method were estimated to adjust for differences between library sizes^36^. Each raw read count was multiplied by the product of the corresponding gene-specific normalization factor and the TMM scale factor to obtain a normalized count. The average expression level of each gene was represented by the mean normalized count across all replicates of each genotype in our data set. After estimation of the density distribution, the 0.25 quantile of the non-zero average expression levels was set as the threshold for calling the activity status of each gene in each sample. Thus, a gene was called active if the average expression level across all replicates was greater than the threshold and otherwise inactive for each genotype. Genes active in only one parent but also the hybrid are designated single parent expression (SPE) or SPE complementation genes, as the expression of only one parent is complemented in the hybrid. We identified these, by comparing the activity of each gene in the hybrids and their corresponding parents. From the classification of the IBM-RIL regions, we can deduct the genotypes of the SPE genes in the hybrids. Based on these genotypes, we classified our SPE genes into those within heterozygous (B73/Mo17 in B73×IBM-RILs, Mo17/B73 in Mo17×IBM-RILs) or homozygous regions (B73/B73 in B73×IBM-RILs, Mo17/Mo17 in Mo17×IBM-RILs). We further distinguished the SPE by which parent was active and indicated the active parent in bold. So, for example a SPE gene in a heterozygous region of a B73×IBM-RIL, where the IBM-RIL parent is active, but the B73 is not, the pattern would be B73/**Mo17** (Figure 2A). For a SPE gene in a homozygous region of a Mo17×IBM-RIL, where the Mo17 parent is active, but the IBM-RIL is not, the pattern would be **Mo17**/Mo17 (Figure 2B).

**Figure 2:**
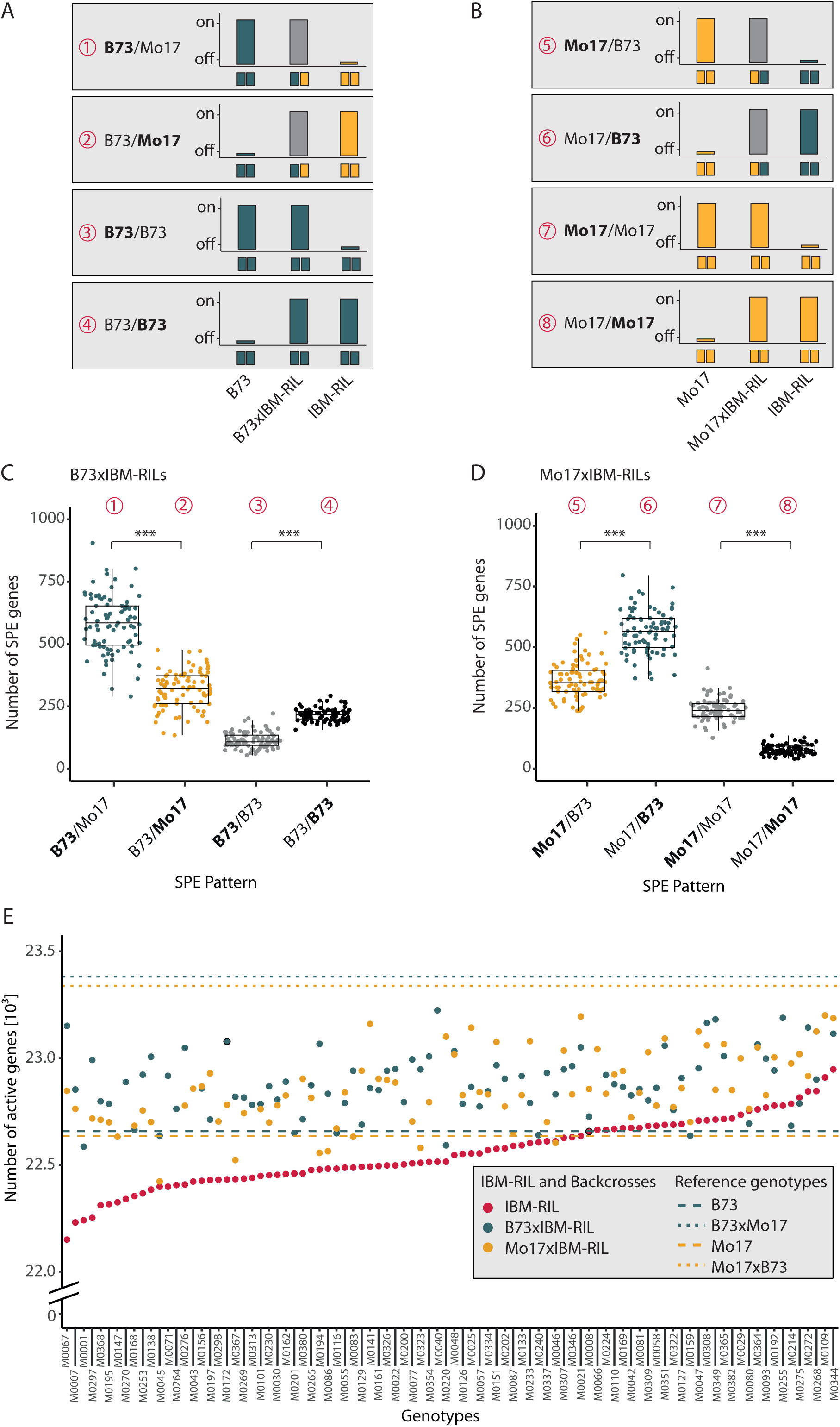
Single-parent expression (SPE) complementation. **A** SPE pattern present in B73xIBM-RIL backcross hybrids relative to the genomic composition of the paternal IBM-RIL and the active parent (Pattern 1-4). **B** SPE pattern in Mo17xIBM-RIL backcross hybrids relative to the genomic composition of the paternal IBM-RIL and the active parent (5-8). A simplified depiction of the activity (on/off) of SPE genes is shown for each SPE pattern above the genotypic composition of the respective pattern for each parent and hybrid. Pattern 1, 2 and 5, 6 are also present in B73xMo17 and Mo17xB73 respectively. **C** Boxplots displaying the total number of SPE genes for each of the possible SPEpattern (1-4) in the B73xIBM-RIL hybrids (N= 85). **D** Boxplots displaying the total number of SPE genes for each of the possible SPE pattern (5-8) in the Mo17xIBM-RIL hybrids (N=82). Asterisks indicate significant differences (α<0.0001, p-values given as <0.0001 in other words, zero) identified by a gaussian mixed model with the hybrid as random effect, the SPE pattern and non-SPE pattern as a fixed factor and a diagonal variance component for the SPE pattern. **E** For each IBM-RIL line (red) and the corresponding B73xIBM-RIL (blue) and Mo17xIBM-RIL (yellow) backcross hybrids the number of active genes is displayed. The dashed lines represent the number of active genes in the inbred lines B73 (blue) and Mo17 (yellow). The dotted lines represent the number of active genes in the reciprocal hybrids B73xMo17 (blue) and Mo17xB73 (yellow).

### Proportion of heterotic variance explained by the number of SPE genes underlying mid-parent heterosis (MPH) of root phenotypes

To estimate the fraction of the heterotic variance explained by the number of genes displaying SPE patterns, we propose the parameter 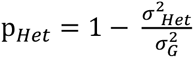, where 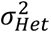 defines the total genetic variance across the hybrid genotypes and 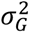 is the genetic variance of the heterosis effect not associated with the number of SPE genes^37,38^

For each backcross population, the genetic variance of the mid-parent heterosis (MPH) effects 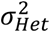 was estimated separately in a “full” regression model (2) based on an extension of the baseline model (1). For this purpose, we defined for each parental genotype (i.e. each the IBM-RIL and the two common parental inbred lines B73 and Mo17) covariates (*x_i_*_1−*x*_). These covariates were initially all set to 0 for each observation. For observations on the parental genotypes, the corresponding covariate for that specific parent was set to 1. For the observation on the hybrids, the two covariates corresponding to its two parents were set to 0.5. Thus, collectively these *x_i_*_1−*x*_ covariates model the effect of the *per se* performance of the parents and the mid-parent values of the hybrids.

MPH was modelled by a regression on the number of SPE genes. For this purpose, the number of SPE genes was set to 0 for all parental genotypes. This was done to be able to include them in the overall model. However, the parental genotypes have no impact on the regression, because their effect is fully absorbed by the covariates for the parental genotype effects.

As the MPH effects of the hybrids were not expected to fall on the regression line, we allowed for deviations from the regression by adding a random effect for hybrids. This was implemented by fitting the random effect z*genotype, where z is a continuous dummy covariable with z=0 for the parental genotypes and z=1 for the hybrids. This dummy variable acts as a switch that turns the random effect off for parental genotypes and on for hybrids ^39^.

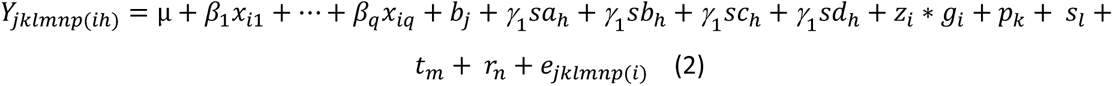

*Y_jklmnp_*_(*ih*)_ represents the parental effect value contributing to MPH of the corresponding hybrid *h* of a specific trait of interest. *µ* represents the intercept, *x_iq_* represents the parental covariables of parent *q* for genotype *i*, *b_j_* represents the fixed effect for block *j*, *sa_h_*, *sb_h_*, *sc_h_* and *sd_h_* represent the covariables for the number of genes displaying pattern 1-4 (Figure 2A) or 5-8 (Figure 2B) for hybrid *h*, respectively. *z_i_* * *g_i_* represents the random effect for hybrid *h* (corresponding to genotype *i*), *z_i_* is a dummy variable with *z_i_* = 0 for parents and *z_i_* = 1 for hybrids^39^. Variable *p_k_* represents the random effect for batch *k* nested within block *j*, *s_l_* represents the random effect for system *l* nested within batch *k* and block *j*, *t_m_* represents the random effect for triplet *m* nested within system *l*, batch *k* and block *j*, *r_n_* represents the random effect for row *n* nested within triplet *m*, system *l*, batch *k* and block *j*, and *e_jklmnp_*_(*i*)_ represents the random error effect for plant *p* of genotype *i* in block *j*, batch *k*, system *l*, triplet *m* and row *n*.

The analysis was implemented in R (version 4.0.1) using the lme4 package (version 1.1-29). In contrast to the baseline model, the fixed effect for genotype was replaced by individually defined covariates of the parental genotypes and fixed effects for the number of SPE genes.

To determine 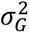 a “null” model (3) excluding the fixed effects of the covariates accounting for the number of SPE genes, was fitted.

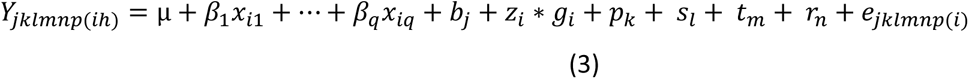

where the notations are the same as in model (2).

### Expression Quantitative Trait Loci (eQTL) analysis

An eQTL analysis was performed with the R/qtl2 package (version 0.22)^40^ to identify positions that were significantly associated with gene expression values based on the masked and filtered SNP data. For each of the three cross-types (IBM-RIL, B73×RILs, Mo17×RILs) the classified and filtered SNP loci within B73 or Mo17 regions in the IBM-RILs were taken as marker input data. The positions of these SNP loci were used as preliminary genomic positions, as well as physical positions. The estimated expression means obtained from the model coefficients within the differential expression analysis of each genotype and gene were used as the phenotype data input in R/qtl2. As additional specifications to write the control file, the cross type was set to “rilself” for RILs by selfing, the alleles were set to “A” and “B” for B73 and Mo17 and the genotype codes were set to A/A=1 and B/B=2 to specify the transformation of homozygous alleles into numeric values (https://github.com/agroot-ibed/r-qtl2-analysis, updated on 12/19/2023, checked on 12/19/2023). Samples with more than 19% missing genotypes were dropped, as well as duplicated genotypes and markers with more than 60% missing genotype information. The genetic map was estimated from the physical positions and genotype information by the est_map() function with parameters maxit = 2000, error_prob = 0.001 and tol = 0.0001. The reduce_markers() function was used to retain only markers that were ≥1 centiMorgan (cM) apart to avoid retaining an excess of redundant markers. Pseudomarkers were inserted at a distance of 1 cM to the existing markers. A hidden Markov model calculated the genotype probabilities at all positions, with error_prob = 0.001. This was followed by a genome scan, which was done by a Haley-Knott regression^41^ to establish the association between genotype and expression phenotype with a linear model. In simple words, within each eQTL analysis, each marker is tested to see whether there is an association with the expression of single genes, the result is an LOD curve. In order to find out whether the highest LOD value is significant, a permutation was carried out and all significant peaks were saved. In more detail, to calculate the adjusted *p*-values for the resulting logarithm of odds (LOD) scores for a single gene, 10 000 permutations were done, reshuffling the expression data randomly and recording the maximum LOD score of each permutation. Selecting a significance threshold of α ≤0.001, we used the 99.9^th^ percentile of the ordered LOD maxima as the threshold to detect a significant eQTL for the gene^42^. The genomic map was converted to a physical map with the interp_map() function. By selecting the respective threshold, the physical position, confidence interval and LOD of significant peaks was obtained by the find_peaks() function^40^. The exact adjusted *p*-values were determined by calculating the percentile of permutation maxima, higher than the respective LOD^42^. This process was repeated for all (37782) active genes. To subsequently also correct for the testing of multiple genes, the false discovery rate was used on the adjusted *p*-values of the LOD peaks of all genes with p.adjust() function setting method to “FDR” and n to the total number of genes plus the number of second and third significant peaks. Peaks with an FDR ≤ 0.001 were considered significant. Start and end position of genes were added from the annotation file. The same procedure was performed on all three cross-type data sets (IBM-RIL, B73×IBM-RIL, Mo17×IBM-RIL). The resulting eQTL peaks were combined and distinct eQTLs were selected: in cases where multiple eQTLs were identified for a gene, we assessed whether the different peak positions corresponded to different regulatory elements. If eQTLs for the same gene were ≥25 Mbp apart or on different chromosomes and their positions did not lie within the confidence intervals of each other, they were considered to be different from each other and were retained. If multiple eQTLs for the same gene did not differ by the specified standards but were in close proximity to each other, only the eQTL with the shortest confidence interval or the highest LOD in case of equal confidence intervals was retained in the merged list. The eQTLs were classified into *cis* and *trans* eQTLs based on their distance from the start of their respective gene. *Trans-*regulating eQTLs were defined as located at a distance of at least 2.5 Mbp from the start of the gene and where their confidence interval did not include the start of the gene. *Cis-*regulating eQTLs were defined as located in proximity to the start of the gene (<2.5 Mbp) or located such that their confidence interval includes the start of the gene (Data file S4).

### Transcriptome Wide Association Analysis (TWAS)

A TWAS was conducted to associate gene expression levels of active genes with phenotypic traits. The active genes were filtered to select those genes which were active in at least 5% of all genotypes (14). We used the MLMM^43^, BLINK^44^ and FarmCPU^45^ models, implemented in the R package GAPIT (version 3)^46^, including the three first principal components for the initial identification of genes and in case of the MLMM the variance-covariance matrix between individuals as kinship. Each population (IBM-RILs, B73×IBM-RILs, Mo17×IBM-RILs) was analyzed separately. For use in GAPIT, expression values were rescaled to values between 0 and 2 for each population. The presented TWAS + SPE candidate genes (TSG) were additionally investigated using Student’s t-test.

### Determination of syntenic and non-syntenic genes

The syntenic and non-syntenic genes were determined by comparison against a published list of syntenic grass genes^47^. Those genes with *cis*-regulatory eQTL were compared against those with *trans*-regulatory eQTL using the Fishers’ exact test in R with fisher.test().

## Results

### Transcriptome profiling and sample evaluation via SNP calling reveals cross-over locations and IBM-RIL specific regions

After seed propagation, we selected 112 IBM-RILs and their B73×IBM-RIL and Mo17×IBM-RIL hybrids based on seed availability together with the reciprocal hybrids Mo17×B73 and B73×Mo17 as well as the parents B73 and Mo17 for subsequent phenotyping and RNA-sequencing of young primary roots. After SNP calling and quality control (see methods), we classified the IBM-RIL genomes into B73 and Mo17 regions. During this process, we masked IBM-RIL specific regions which contained SNPs not present in Mo17 or B73 and are thus likely the result of contamination from other genotypes. In total, 834 out of 1152 sequenced samples passed all quality thresholds and were subjected to further analyses. They consist of 2-3 biological replicates of 85 B73×IBM-RIL and 82 Mo17×IBM-RIL backcross hybrids and their IBM-RIL parents together with 23 (B73×Mo17) and 24 (Mo17×B73) biological replicates of the full reference hybrids and 47 biological replicates of B73 and 42 of Mo17 (Data file S1). The categorization of the IBM-RIL genomes revealed the putative recombination breakpoints between Mo17 and B73 regions (Figure S2).

We explored the sample quality and the relationship among the 834 RNA-seq samples in a multidimensional scaling (MDS) plot (Figure 1C). The different genotypes (B73, Mo17, B73×Mo17, Mo17×B73) and populations (IBM-RILs, B73×IBM-RILs, Mo17×IBM-RILs) form distinct clusters, which are clearly separated from each other. As expected, the clusters of the two reciprocal reference hybrids B73×Mo17 and Mo17×B73 overlap because their genomes are identical. These hybrid samples are located in between their parental inbred lines on dimension 1 of the MDS plot (Figure 1C). Similarly, the IBM-RILs are located between their original parental inbred lines B73 and Mo17. Finally, as expected the two backcross populations B73×IBM-RIL and Mo17×IBM-RIL are located between their parental lines (Figure 1C).

### Root traits display heterosis in backcross populations

For each parent-hybrid combination, we determined lateral root density (Figure S3A), total number of root tips (Figure S3B), total root length (Figure S3C) and total root volume (Figure S3D). The inbred line B73 outperformed the inbred line Mo17 in all measured root traits. The B73×Mo17 hybrid displayed higher values than the Mo17×B73 hybrid for the total number of root tips, total root length and total root volume. In contrast, Mo17×B73 displayed a higher lateral root density than B73×Mo17 (Figure S3).

In addition, we estimated mid-parent heterosis (MPH) for all measured root traits. The level of MPH of the fully heterozygous B73×Mo17 and Mo17×B73 hybrids (from 49% for lateral root density to 127% for the number of root tips) exceeded the average of the partially heterozygous B73×IBM-RIL and Mo17×IBM-RIL hybrids (from 25% for lateral root density to 62% for total root volume) in all traits. Nevertheless, most of the IBM-RIL backcross hybrids displayed substantial MPH (Figure S3, Data file S5).

### Heterozygosity promotes single parent expression complementation

Genes which are active in one parental inbred line and in the F_1_ hybrid offspring, but inactive in the second parent, display single-parent expression (SPE). In the highly heterozygous reference hybrid B73×Mo17 we identified 1297 SPE **B73**/Mo17 genes, active in the hybrid and the maternal inbred line B73 (pattern 1, Figure 2A; allele contributed by active parent in bold). In the reciprocal hybrid Mo17×B73 we identified 1241 SPE **Mo17**/B73 genes active in the hybrid and only in the maternal inbred line Mo17 (pattern 5, Figure 2B). In addition, we identified 1228 SPE B73/**Mo17** (pattern 2) and 1253 SPE Mo17/**B73** (pattern 6) genes active in the paternal inbred line and the hybrid, but not the corresponding maternal parent. Between both reference hybrids, 85% of the SPE genes were conserved (Data file S6).

Within the two IBM-RIL backcross populations we identified eight different SPE patterns, four in each backcross population, depending on the genomic composition of the gene in the paternal IBM-RIL. SPE genes of all patterns are expressed in the hybrid. But we can distinguish them by the active parent (Figure 2A, B; allele contributed by active parent in bold). Pattern 1 (B73×IBM-RILs): Genes lie within heterozygous regions of the hybrid and the expressed parent is B73; Pattern 2 (B73×IBM-RILs): Genes lie within heterozygous regions of the hybrid and the expressed parent is the IBM-RIL, which contributes the Mo17 allele for these genes; Pattern 3 (B73×IBM-RILs): Genes lie within homozygous regions of the hybrid and the expressed parent is B73; Pattern 4 (B73×IBM-RILs): Genes lie within homozygous regions of the hybrid and the expressed parent is the IBM-RIL, which contributes the B73 allele for these genes; Pattern 5 (Mo17×IBM-RILs): Genes lie within heterozygous regions of the hybrid and the expressed parent is Mo17; Pattern 6 (Mo17×IBM-RILs): Genes lie within heterozygous regions of the hybrid and the expressed parent is the IBM-RIL, which contributes the B73 allele for these genes; Pattern 7 (Mo17×IBM-RILs): Genes lie within homozygous regions of the hybrid and the expressed parent is Mo17; and Pattern 8 (Mo17×IBM-RILs): Genes lie within homozygous regions of the hybrid and the expressed parent is the IBM-RIL, which contributes the Mo17 allele for these genes. (Figure 2A, B).

We observed in the B73×IBM-RIL and Mo17×IBM-RIL backcrosses on average 1229 (Figure 2C) and 1247 (Figure 2D) SPE genes per hybrid, which is ~50% of the number of SPE genes in B73×Mo17 and Mo17×B73. Across B73×IBM-RIL and Mo17×IBM-RIL backcross hybrids, SPE patterns in heterozygous regions (Figure 2C, D: patterns 1, 2, 5, 6) occurred more often than in homozygous regions (Figure 2C, D: patterns 3, 4, 7, 8). We identified more SPE genes in heterozygous regions with B73 as the active parent (Figure 2C, D: pattern 1 and 6) than with Mo17 as the active parent (Figure 2C, D: pattern 2 and 5). In regions homozygous for B73, more genes with paternal activity were observed (Figure 2C: patterns 4 vs 3), whereas in regions homozygous for Mo17 more genes with maternal activity were determined (Figure 2D: pattern 7 vs 8).

As a consequence of expression complementation of genes expressed in only one parent, which are then also active in the hybrid (SPE), we observed that hybrids express more genes than their parental lines (Figure 2E). While the inbred lines B73 and Mo17 express 22 658 and 22 635 genes, respectively, their reciprocal F_1_ hybrid offspring B73×Mo17 and Mo17×B73 display 736 and 692 more active genes than their parental average. The corresponding B73×IBM-RIL and Mo17×IBM-RIL backcrosses expressed on average 278 and 253 more genes than their parents. Furthermore, the number of active genes in the backcross hybrids is positively correlated with the fraction of heterozygous genomic regions in the hybrid (Figure S5). These results indicate that the level of heterozygosity is important for expression complementation in hybrids.

### Contribution of SPE to heterosis

To estimate the proportion of heterosis variance explained by the number of SPE genes underlying MPH of root traits, we determined in each backcross population the coefficient of determination p_HET_. Across the four examined root traits, 12% (total root volume) to 29% (total number of root tips) of the heterotic variance was explained by the number of SPE genes across the four different patterns in the B73×IBM-RIL backcross hybrids. In contrast, in the Mo17×IBM-RILs only between 4% (lateral root density) and 9% (total root volume) of the heterotic variance was explained by the total number of SPE genes. For the traits of total number of root tips and total root length in the Mo17×IBM-RILs we observed a negative proportion (−8% and −0.5%) of explained heterotic variance of the number of SPE genes on MPH (Table 1).

**Table 1:** Proportion of heterotic variance explained by the number of SPE genes on mid-parent heterosis for different root phenotypes.

| Trait | B73xIBM-RILs |  |  | Mo17xIBM-RILs |  |  |
| --- | --- | --- | --- | --- | --- | --- |
| | $\sigma^2_{\text{Het}}$ | $\sigma^2_{\text{G}}$ | $\rho_{\text{Het}}$ | $\sigma^2_{\text{Het}}$ | $\sigma^2_{\text{G}}$ | $\rho_{\text{Het}}$ |
| **No. Of root tips** | 0.232 | 0.325 | 0.29 (29%) | 0.205 | 0.190 | -0.08 (-8%) |
| **Total root volume** | 2.366 | 2.701 | 0.12 (12%) | 1.022 | 1.121 | 0.09 (9%) |
| **Total root length** | 2.715 | 3.561 | 0.24 (24%) | 2.202 | 2.191 | -0.01 (-1%) |
| **Lateral root density** | 5.461 | 6.805 | 0.20 (20%) | 6.675 | 6.925 | 0.04 (4%) |

$\sigma^2_{\text{Het}}$ = unexplained genetic variance of heterosis effect, not associated with SPE genes; $\sigma^2_{\text{G}}$ = total genetic variance among the hybrid genotypes; $\rho_{\text{Het}}$ = Coefficient of determination: proportion of the heterotic variance explained by the number of SPE genes.

### *Trans*-regulatory elements are more frequent in Mo17 than B73

An expression quantitative trait locus (eQTL) is a position in the genome that is significantly associated with expression variation of a gene and thus likely regulates this gene. We identified 13,778 eQTL for 13,434 protein-coding genes. While most genes were regulated by a single eQTL, 334 genes were regulated by two eQTL and five by three eQTL (Data S4). We categorized all eQTL into either *cis-* or *trans*-regulating by their location relative to the starting position of the corresponding gene. We defined *cis*-regulating eQTL as located at a distance of <2.5 Mbp from the start codon of their target gene on the same chromosome. In contrast, most *trans*-regulating eQTL were located on chromosomes other than those of their target gene (83%). The remaining *trans* eQTL were located on the same chromosome as their target at a distance of ≥2.5 Mbp (and the target gene was outside of the eQTL confidence interval). The logarithm of odds (LOD) is the significance measure of an eQTL being present using the Haley-Knott regression. The median LOD was 15.5 for *trans*- and 16.3 for *cis*-eQTL (Data S4). Effect sizes usually estimate how big the effect of each identified locus is on the phenotype, or in this case on gene expression. As LOD values and effect size estimates of the eQTL on the gene expression are correlated^48^, this indicates similar effect sizes of *cis-* and *trans*-acting eQTL. In total, 88% (12,057/13,778) of all identified eQTL acted in *cis*, while 12% (1,701/13,778) acted in *trans* on their target genes. For genes active in the B73 reference genotype 7% (771) of their eQTL were acting in *trans* while for genes active in the Mo17 reference genotype almost twice as many eQTL acting in *trans* were identified (13%, 1,439). This suggests that Mo17 contains more *trans*-acting eQTL than B73. The values for all other genotypes show *trans*-acting eQTL ratios between the B73 and Mo17 values (Table 2).

**Table 2:** *Trans* eQTL for active genes.

| Genotype | *Trans*-regulating eQTL [in % of eQTL for active genes] | Number of *trans*-regulated genes |
| --- | --- | --- |
| B73 | 7 | 771 |
| B73_Mo17 | 11 | 1365 |
| Mo17 | 13 | 1439 |
| Mo17_B73 | 11 | 1363 |
| IBM-RILs* | 10 | 1107 |
| B73xIBM-RILs* | 9 | 1066 |
| Mo17xIBM-RILs* | 12 | 1385 |

\*Values are the mean of the respective category

### Heterozygous SPE genes with Mo17 activity are *trans*-regulated disproportionately more often

In both fully heterozygous reference hybrids B73×Mo17 and Mo17×B73, we detected eQTL for 85% of the SPE genes (Data file S6). Overall, 95% of eQTL for SPE genes with B73 contributing the active parental allele were *cis*-regulating (Figure S6). By contrast, SPE genes with Mo17 as the active allele were predominantly (58 - 59%) *trans*-regulated (Figure S6).

Similarly, in B73×IBM-RIL and Mo17×IBM-RIL backcross hybrids (Figure S7), heterozygous SPE genes where B73 is the active allele are regulated almost exclusively by *cis* eQTL (on average 95% and 96%) (Figure S7A: pattern 1; Figure S7B: pattern 6). By contrast, on average only 60% (Figure S7A pattern 2) and 62% (Figure S7B pattern 5) of heterozygous SPE genes with an active Mo17 allele are *cis*-regulated. Hence, in general heterozygous SPE genes with an active Mo17 allele are significantly more frequently *trans*-regulated than those with B73 as the active parental allele (Table 2).

### Homozygous SPE genes are partially *trans*-regulated

SPE genes located in homozygous regions showed different ratios of *cis-* to *trans-*eQTL regulation, based on the active parent. While in homozygous B73 regions (occurring in B73×IBM-RILs), SPE with maternally active alleles (**B73**/B73, Figure S7A, pattern 3) were primarily *cis*-regulated (87%), those with paternally active alleles (B73/**B73,** Figure S7A pattern 4) were primarily *trans*-regulated (67%). In homozygous Mo17 regions (Mo17×IBM-RILs) we saw the opposite. SPE genes with maternally active alleles (**Mo17**/Mo17, Figure S7B pattern 7) showed more *trans*-regulating eQTL (58%) and for SPE with paternal activity (Mo17/**Mo17,** Figure S7B pattern 8) the majority of eQTL (81%) were *cis*-regulating. Interestingly, the SPE patterns 4 and 7, which are primarily regulated by *trans*-eQTL, also had higher absolute numbers of SPE genes (Figure 2A, B). The observed proportions of *trans*-regulation for SPE patterns 1 to 8 were significantly different from the ratios in non-SPE genes, of which 91% were *cis*-regulated (Figure S7).

### SPE genes are predominantly regulated in heterozygous regions

Overall, we observed that for heterozygous SPE genes, both *cis*- and *trans*-regulating eQTL are predominantly located in heterozygous genomic regions (Figure 3). We observed that *cis*-regulating eQTL located in hetero- and homozygous regions regulate SPE genes which are also hetero- or homozygous as the corresponding eQTL. This is expected, as *cis*-eQTL are located in close proximity to their gene (Figure 3A, Figure 3B). In contrast, target genes of *trans*-regulating eQTL are randomly distributed across the genome. Accordingly, *trans*-acting eQTL regulate SPEs in both homozygous and heterozygous regions with equal frequency, independent of whether the eQTL is homo- or heterozygous (Figure 3A, Figure 3B). Together with the association of Mo17 activity with *trans*-regulation, this leads to the observed ratios of *cis*- to *trans*-regulation in homozygous SPE patterns. For instance, eQTL regulating homozygous genes in *trans* (Figure 3A and 3B, patterns 4 and 7) were located almost exclusively in heterozygous regions. In both cases, the active parent carries the Mo17 allele at the eQTL position which is responsible for the gene activity of those SPE genes (Figure S8 A, B). This further explains the higher proportion of *trans*-regulation of pattern 4 and 7 as well as the generally higher number of SPE genes for these patterns compared to pattern 3 and 8. In summary, SPE patterns with parental B73 activity showed a slightly higher proportion of regulation by *cis*-acting eQTL compared with non-SPE genes, while the SPE patterns with Mo17 activity showed substantially lower regulation by *cis-*acting eQTL and thus higher *trans*-regulation (Figure 3, Figure S7).

**Figure 3:**
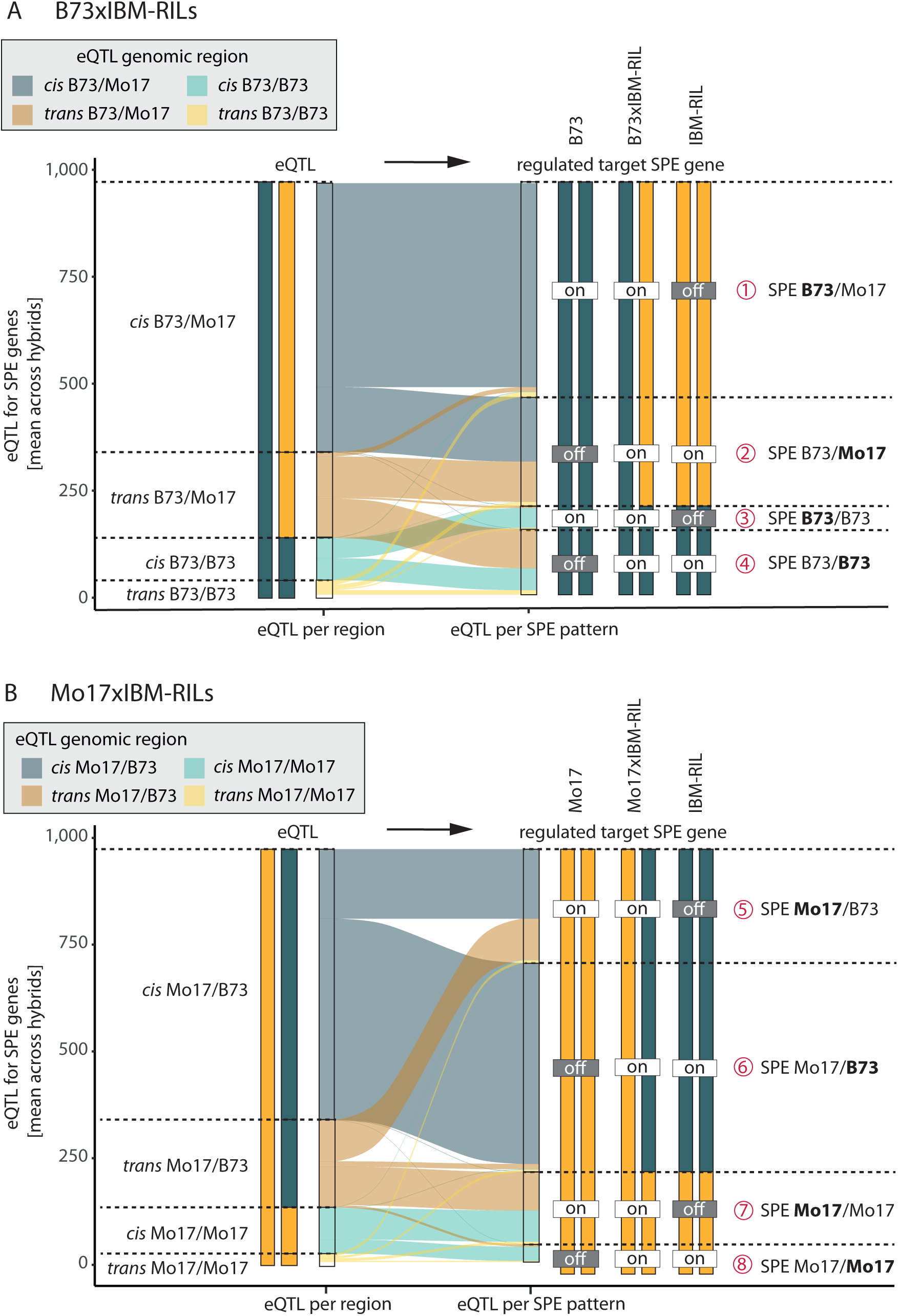
The average number of eQTL per genomic region for the different SPE pattern in **A** B73xIBM-RILs and **B** Mo17xIBM-RILs is shown. The width of the connecting bands corresponds to the average number of eQTL. The shade indicates the genomic region of the eQTL, (dark = heterozygous, light = homozygous) and blue color corresponds to *cis* and orange to *trans*-regulation. The genotypes at the eQTL position and regulated SPE genes are indicated as bars (yellow = Mo17 allele, truqoise = B73 allele), and the gene activity for SPE is shown as (on/off).

### *Trans*-regulated genes and SPE genes are often non-syntenic

The maize genome contains genes with orthologs of positional synteny in sorghum, suggesting these genes are highly conserved across evolution, and a set of genes without syntenic partners (non-syntenic) in sorghum and other grass species, which are therefore most likely evolutionarily younger or changed their position during evolution^49^. In the B73v5 maize genome, 40% (15 612) of genes can be classified as non-syntenic. Among the active genes in this study, 27% (7622) were non-syntenic. By contrast, on average 58% of SPE genes in the hybrids are non-syntenic.

Interestingly, genes with *trans*-eQTL were more likely to be non-syntenic (70%) than genes with *cis*-eQTL (22%) across all expressed genes (Table S1). Both *cis-* and *trans*-regulated SPE genes were more frequently non-syntenic than non-SPE genes (Figure 4). Additionally, non-syntenic genes were particularly prominent among the t*rans*-regulated SPE genes with Mo17 activity. For example, 96% of *trans*-regulated pattern 2 genes (B73/**Mo17**) were non-syntenic (Figure 4). It should be noted that these are relative values: the absolute numbers of genes in patterns 3 and 8 are low in general, and patterns 1 and 6 are not often *trans*-regulated (Figure 3).

**Figure 4:**
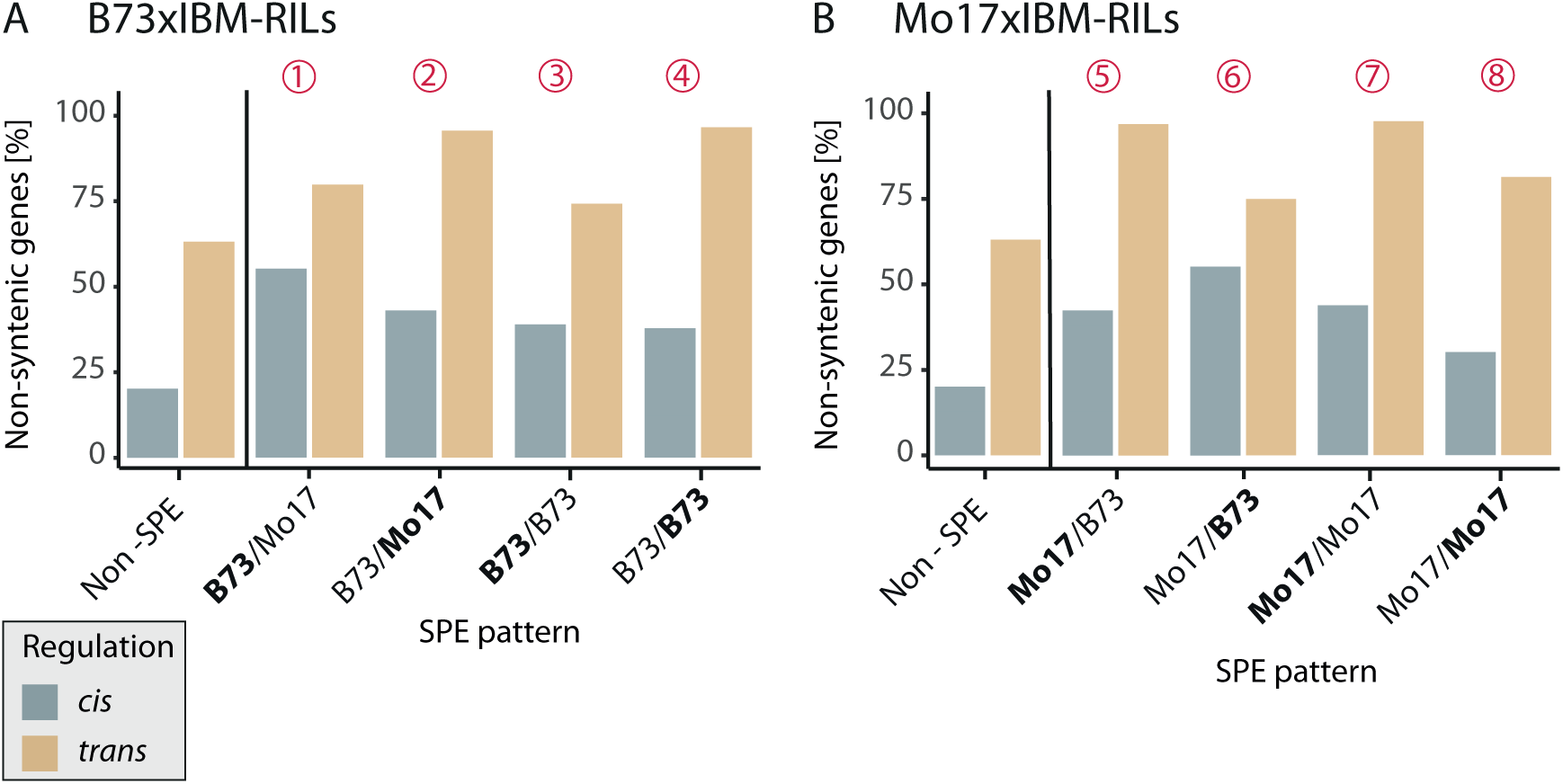
Synteny of SPE pattern genes. The average proportion of non-syntenic genes (vs. syntenic genes) among *cis*- (blue) and *trans* (yellow)-regulated SPE genes. Percentages are an average of genes in the respective category across all (**A**) B73xIBM-RILs (N=85) or (**B**) Mo17xIBM-RILs (N=82).

### SPE gene expression influences lateral root density in different ways

We performed a transcriptome wide association analysis (TWAS) to identify genes whose expression values associate with phenotypic traits. We analyzed four root traits (lateral root density, number of root tips, total root length, total root volume) and the two IBM-RIL backcross hybrid populations B73×IBM-RIL and Mo17×IBM-RIL separately, as different mechanisms of expression regulation and expression levels might be observed between the populations. We found 18 positively and 17 negatively correlated genes across the two populations. Only one gene was identified in both populations (Data file S7). Therefore, different genes might control hybrid vigor of roots in the different populations.

Among the 35 TWAS genes whose expression correlated with phenotypic traits, 7 showed SPE complementation in more than 10 hybrids. We designated these TWAS & SPE (TSG) genes (Data file S8). All of the TSG were identified in the B73×IBM-RIL hybrid population. For lateral root density, 10 TWAS genes were identified by the BLINK and MLMM models of GAPIT in total, and 4 of these genes also showed an SPE pattern (TSG) (Data file S8, Figure 5A). We followed up on the two TSG which showed an SPE pattern in most hybrids (TSG 1 & TSG 2) (Figure 5B, C). TSG 1 (Zm00001eb349930) showed an SPE pattern in 30 of the 85 B73×IBM-RIL hybrids (Figure 5B, Data file S8), predominantly the SPE B73/Mo17. The gene was cis-regulated and non-syntenic. B73×IBM-RIL hybrids with an active TSG 1 showing an SPE pattern had significantly lower lateral root density (α ≤0.05, Figure 5B).

**Figure 5:**
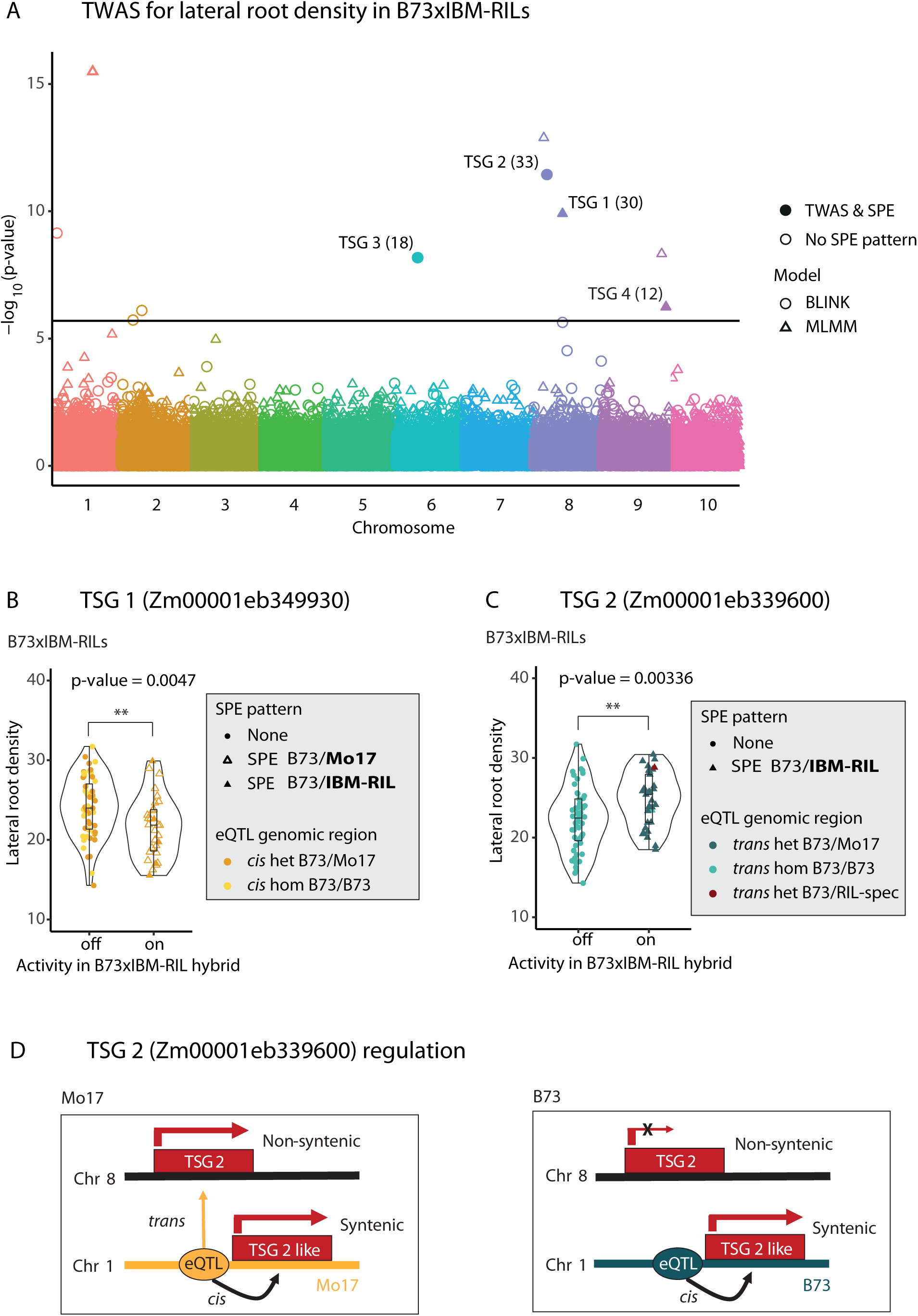
Transcriptome Wide Association Analysis (TWAS). **A** Manhattan plot of TWAS analysis for lateral root density in B73xIBM-RIL hybrids. Each point shows a gene with the p-value of the association between the genes expression with lateral root density. Results for the analysis with the BLINK model are shown as circles, results of the MLMM model are shown as triangles. Filled points indicate, significant TWAS genes, that also show SPE pattern (TSG), with the number of hybrids where the gene shows an SPE in brackets. The horizontal line indicates the fdr = 0.05 as per bonferroni correction. **B, C** P-values correspond to Student’s t-test computed independently from the TWAS analysis based on the gene activity. The plots show the lateral root density of each B73xIBM-RIL with either inactive (off) or active (on) **B** TSG 1 Zm0001eb349930 or **C** TSG 2 Zm0001eb339600 in the hybrids. The shape of the points corresponds to the SPE pattern and the color to the genotype at the eQTL position (yellow = *cis* regulating, blue = *trans* regulating, darker color = heterozygous regions, light color = homozygous regions). The color shows the genotype of the hybrid at the eQTL position. N(B73xIBM-RILs) = 85, N(Mo17xIBM-RILs) = 82. **D** shows a model of gene expression of the non-syntenic gene TSG 2 and the syntenic paralog of TSG 2, TSG 2 like, which are regulated from the same eQTL. Depiction shows activity in Mo17 and B73 inbred lines. In heterozygous genotypes, TSG 2 is active, when at least one Mo17 allele is present at the eQTL position. TSG 2 like is active in all genotypes.

A different effect was visible for TSG 2 (Zm00001eb339600), which showed B73/**IBM-RIL** pattern in 33 B73×IBM-RIL hybrids (Figure 5C Data file S8). As there were no SNPs near the gene to classify the paternal allele of the gene itself, the SPE pattern was called B73/**IBM-RIL**, instead of B73/**Mo17** or B73/**B73**. Interestingly, TSG 2 was *trans*-regulated and the genotype at the eQTL position could be determined. In the B73×IBM-RILs, the hybrids with an active TSG 2 had a significantly (α ≤0.05) higher lateral root density and showed the heterozygous B73/Mo17 genotype at the eQTL position (Figure 5C). When searching for sequences similar to TSG 2 (Zm00001eb339600) in the B73 reference genome, we identified a paralog (Zm00001eb039610) and called it TSG 2-like. TSG 2-like is located close to the eQTL position of TSG 2 (<2.5 Mbp) (Data S9, Figure 5D). While the TWAS gene TSG 2 is non-syntenic and *trans*-regulated, its paralog TSG 2-like at the eQTL position is *cis*-regulated from the same eQTL position and syntenic (Figure 5D). The syntenic TSG 2-like gene is expressed in Mo17 and in B73 and expressed in all genotypes. However, the non-syntenic TSG 2 is only active in Mo17, but not in B73 (Figure 5D) and subsequently only active in those hybrids where at least one Mo17 allele is present at the eQTL position (Figure 5C).

## Discussion

SPE complementation, where the hybrid expresses a gene that is only active in one of its parents, was suggested to contribute to the translation of parental diversity into phenotypic heterosis^10^. This concept is in line with the dominance model of heterosis which assumes that many slightly deleterious alleles of the parents are complemented in the hybrid by dominant alleles^7^. The number of SPE genes identified in this study is similar to previous studies for B73xMo17 and Mo17xB73 seedling roots^11,12,50^. In the IBM-RIL backcross hybrids, we identified on average a substantially lower phenotypic mid-parent heterosis, fewer SPE genes compared to the fully heterozygous hybrids and a lower average heterozygosity. We also observed that SPE genes are mainly located in heterozygous regions of the genome, thus explaining the lower numbers of SPE in the backcross hybrids (Figure 2). Hence, the association of the number of SPE genes with heterosis is not only conditioned by genetic diversity^10^ but also by the degree of heterozygosity. As a result of SPE complementation, the number of active genes increased with heterozygosity (Figure S5).

We demonstrated that the number of genes showing SPE complementation in homozygous and heterozygous regions of the genome (Figure 2A-D) explains up to 29% of the heterotic variance in the phenotype of the hybrids (Table 1). As different genes are likely responsible for heterosis in different traits^51^, our finding suggests that SPE as a group of genes are in general responsible for heterosis, but to a different extent for different traits. It is for instance likely that SPE genes influence heterosis in B73xIBM-RILs but to a smaller degree in Mo17xIBM-RILs, which in general generate less vigorous hybrids.

In our study we detected different ratios of *cis* and *trans* regulation among active genes in inbred lines (Mo17:13% trans-regulation; B73; 7% trans-regulation; Table 2). In the reciprocal hybrids, this difference was significantly amplified for SPE genes (Mo17 active:**~**60% *trans*; B73 active:**~**5% *trans*; Figure S6), similar to the IBM-RIL backcross populations (Figure S7). Previously, contrasting *cis-* and *trans-*regulation was associated with parental alleles in another eQTL study, using IBM-RIL backcrosses from a smaller subset of the IBM-RIL population^14^. In that study, 86% of *trans*-regulation was associated with paternal dominance, where the expression level of the paternal allele was adopted^14^. In our study, investigation of the eQTL positions revealed that most SPE genes are regulated by eQTL located in heterozygous regions. Substantial proportions of SPE genes located in homozygous regions are regulated in *trans*-from eQTL in heterozygous regions, leading to higher numbers of these specific SPE patterns (Figure 3, pattern 4 and 7). We showed that for many *trans*-regulated SPE genes, the presence of the Mo17 allele in the eQTL is required for gene activity in the SPE pattern genes in the hybrid (Figure 3, Figure S8). Thus, we did not observe paternal dominance of *trans*-regulated genes regarding SPE genes, but rather a Mo17 dominance of *trans*-regulated SPE genes.

Our present data confirms findings that non-syntenic genes are enriched among SPE genes^11^ and are correlated with heterosis^52^ (Figure 4). We further expanded this concept by demonstrating that non-syntenic genes are enriched among *trans*-regulated genes. Non-syntenic genes have been associated with disease resistance genes^53^ and were suggested to function in environmental adaptation of plants and help hybrids to cope with abiotic stress^50^. We identified a *trans*-regulated non-syntenic SPE candidate gene TSG 2 (Zm00001eb339600), which controls lateral root density in hybrids, whose expression was induced by the Mo17 allele at the eQTL position (Figure 5B). This gene has a syntenic paralog Zm00001eb039610 (TSG 2-like), which is located close to the eQTL position (<2.5Mbp) (Figure 5C, Data S9). Interestingly, this paralog is *cis*-regulated from the same eQTL position as TSG 2 (Figure 5C) and expressed in B73, Mo17 as well as heterozygous genotypes. We hypothesize that there might be a regulatory connection of the *trans*-regulated non-syntenic gene to the syntenic paralog, in the Mo17 genotype. Regulatory interactions of paralogous genes have been previously reported. For instance, the paralogous genes *rcts* and *rctl* are regulated by the same transcription factor^54^ and the syntenic gene *rtcs* recruited younger non-syntenic genes during seminal root evolution^55^. A regulatory connection of *trans*-regulated non-syntenic genes with their syntenic *cis*-regulated paralogs in the Mo17 genotype could explain the high number of *trans*-regulating eQTL among non-syntenic SPE patterns associated with the Mo17 allele. The regulatory differences (B73: *cis*, Mo17: *trans*) between the parents of a hybrid and their different contributions to SPE pattern might be an aspect of how phylogenetic distance is contributing to heterosis.

Among the TWAS genes whose expression correlated with phenotypic traits, we identified genes in the B73xIBM-RIL population for lateral root density, which displayed SPE in a substantial number of parent-hybrid combinations (Data S8, Figure 5). We surveyed the two candidate genes which displayed SPE in the highest number of parent-hybrid combinations in more detail. They showed significantly different lateral root density, based on the activity or inactivity of the gene in the hybrid, and thus the presence or absence of SPE. For the *cis*-regulated gene TSG 1 (Zm00001eb349930) (Figure 5B), a lower lateral root density was observed upon gene activity. In contrast, gene TSG 2 (Zm00001eb339600) (Figure 5C) is *trans*-regulated and displayed a higher lateral root density, upon gene activity, which is induced by the Mo17 allele at the eQTL position.

Thus, we observe regulatory effects of single genes displaying SPE on lateral root density which might help maize to adapt to changing local environmental conditions such as water availability where disparate lateral root densities are beneficial^56^. In summary, we also demonstrated that the association of the number of SPE genes with heterosis is not only conditioned by genetic diversity but also by the degree of heterozygosity. Additionally, SPE mediated phenotypic heterosis, as well as the regulation of SPE genes in IBM-RIL backcross populations depends on the genetic background of the population.

## Data availability

NCBI Bioproject ID PRJNA923128

## Acknowledgements

We would like to thank KWS for the propagation of the IBM-RIL syn. 4 seeds. We thank Helmut Rehkopf (University of Bonn) for his support in propagating the genetic material for this study and the experimental station “Auf dem Hügel” of the University of Bonn. This work was supported by the Deutsche Forschungsgemeinschaft (DFG) Next Generation Sequencing Competence Network (NGS-CN; project 423957469) grant HO 2249/18-1 to F.H. NGS analyses were carried out at the West German Genome Center (WGGC) site in Bonn. The authors acknowledge access to the bonna cluster hosted by the University of Bonn along with the support provided by its High-Performance Computing & Analytics Lab.

## Author contributions

M. P. and J.A.B. carried out the experiments, conducted the statistical analyses, interpreted the data, and drafted the manuscript. A.S.M, G.L, H.S. gave advice regarding population genetics, TWAS analysis and bioinformatics handling of the data. H.-P.P. provided help with the experimental design for the RNA-seq experiment and helped with the statistical analyses. F.H. conceived and coordinated the study and participated in data interpretation and drafting the manuscript. All authors edited the manuscript and approved the final draft.

