## Supplement table S1 for "Regulation of heterosis-associated gene expression complementation in maize hybrids"

Table S1: Synteny of all active genes with eQTL

| Regulation | Syntenic | Non-syntenic |
|---|---|---|
| **Genes with *cis*-eQTL** | 9292 (78%) | 2694 (22%) |
| **Genes with *trans*-eQTL** | 496 (30%) | 1131 (70%) |
