## Supplement Figures S1 to S7 for "Regulation of heterosis-associated gene expression complementation in maize hybrids"

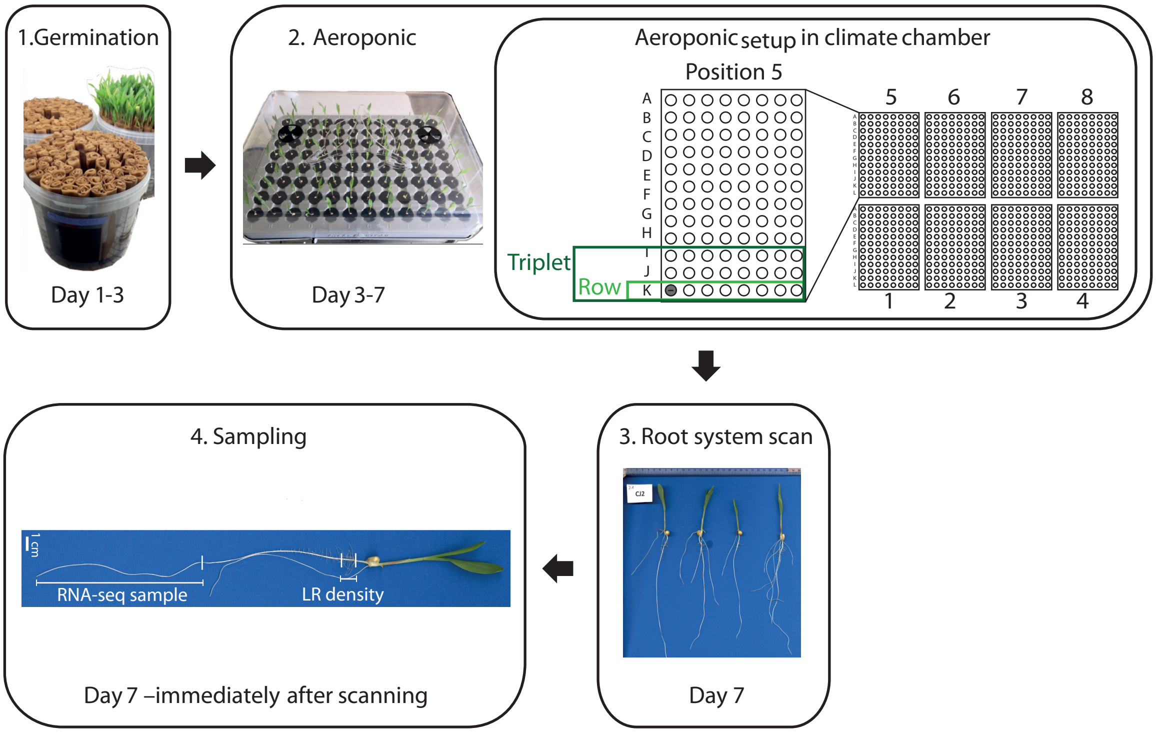


**Figure S1**: Experimental workflow incl. layout of the experimental design. Schematic depiction of the plant growing process from paper rolls until sampling of roots for RNA sequencing. The second box shows the distribution of the aeroponic systems in the climate chamber and the assignment of genotype triplets and individual genotypes within each system.

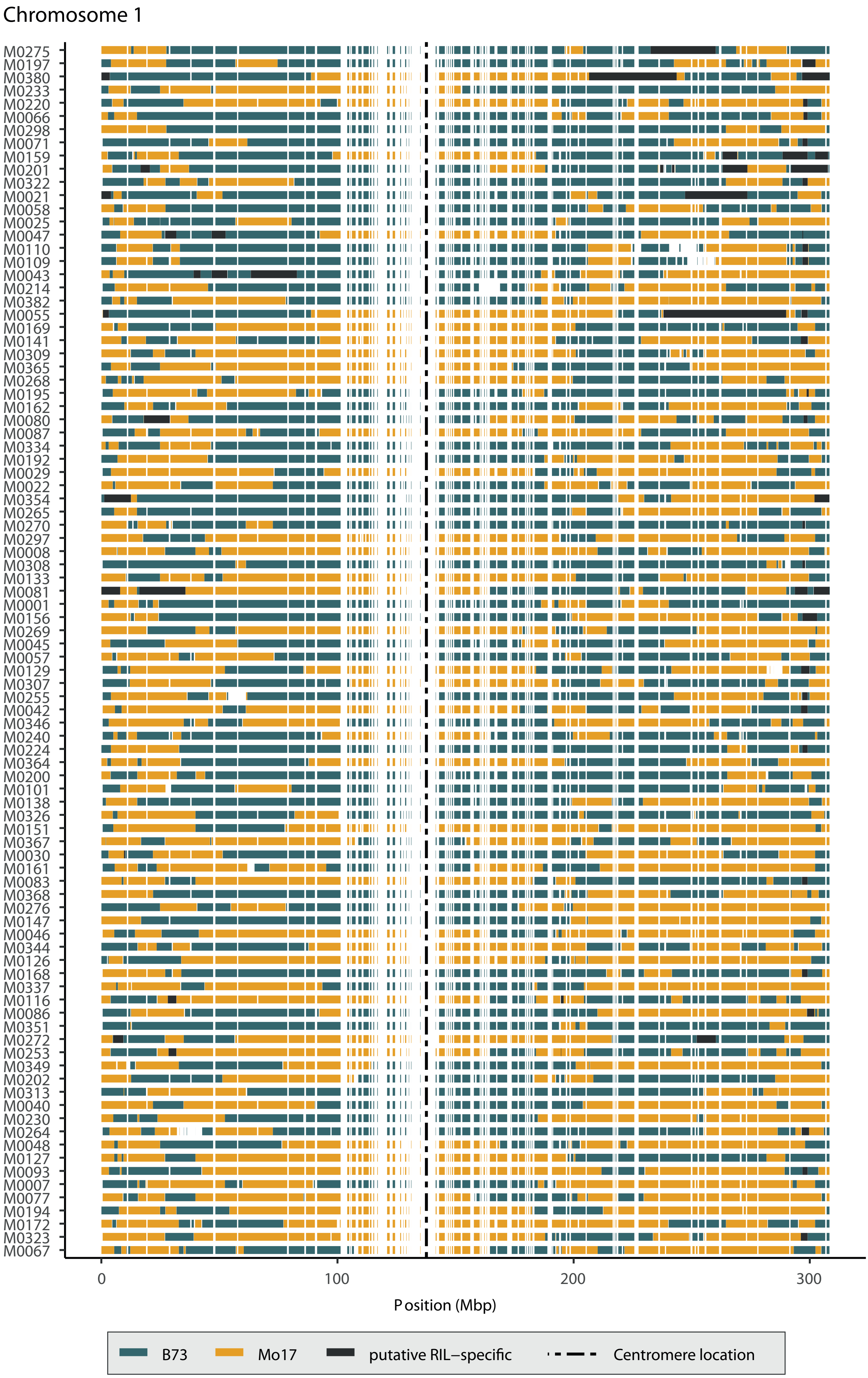


**Figure S2:** Map of genomic regions in IBM-RILs. Each page shows the genomic regions, of all 94 IBM-RILs for one chromosome on physical scale. Regions of B73 are shown in blue, Mo17 in yellow, putative IBM-RIL specific regions, which were masked are shown in black and white spaces indicate, that no SNPs were found in this region. The centromere location is indicated by a vertical dashed line.

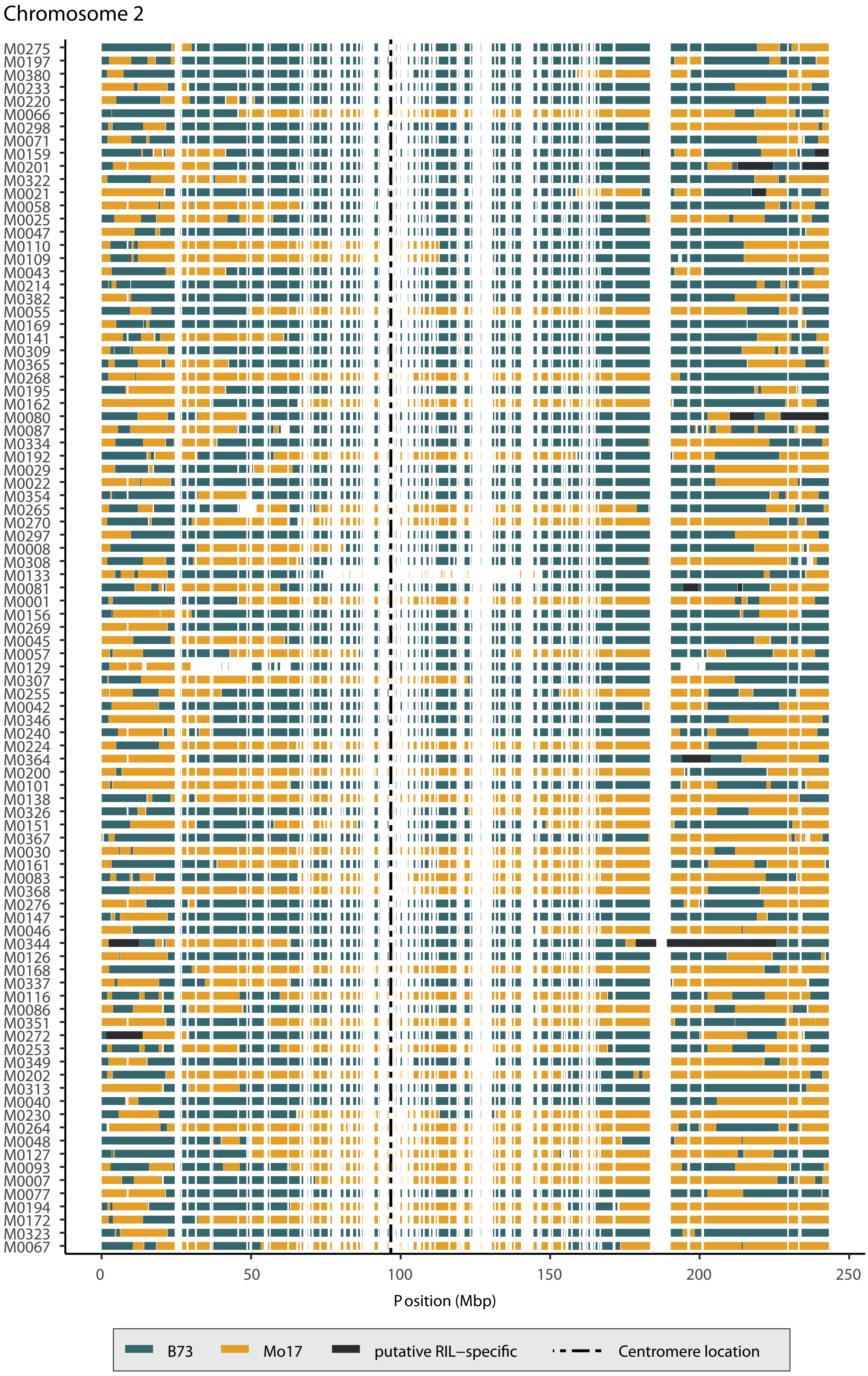
Chromosome 2
M0275
M0197
M0380
M0233
M0220
M0066
M0298
M0071
M0159
M0201
M0322
M0021
M0058
M0025
M0047
M0110
M0109
M0043
M0214
M0382
M0055
M0169
M0141
M0309
M0365
M0268
M0195
M0162
M0080
M0087
M0334
M0192
M0029
M0022
M0354
M0265
M0270
M0297
M0008
M0308
M0133
M0081
M0001
M0156
M0269
M0045
M0057
M0129
M0307
M0255
M0042
M0346
M0240
M0224
M0364
M0200
M0101
M0138
M0326
M0151
M0367
M0030
M0161
M0083
M0368
M0276
M0147
M0046
M0344
M0126
M0168
M0337
M0116
M0086
M0351
M0272
M0253
M0349
M0202
M0313
M0040
M0230
M0264
M0048
M0127
M0093
M0007
M0077
M0194
M0172
M0323
M0067
0
50
100
150
200
250
Position (Mbp)
B73
Mo17
putative RIL−specific
Centromere location

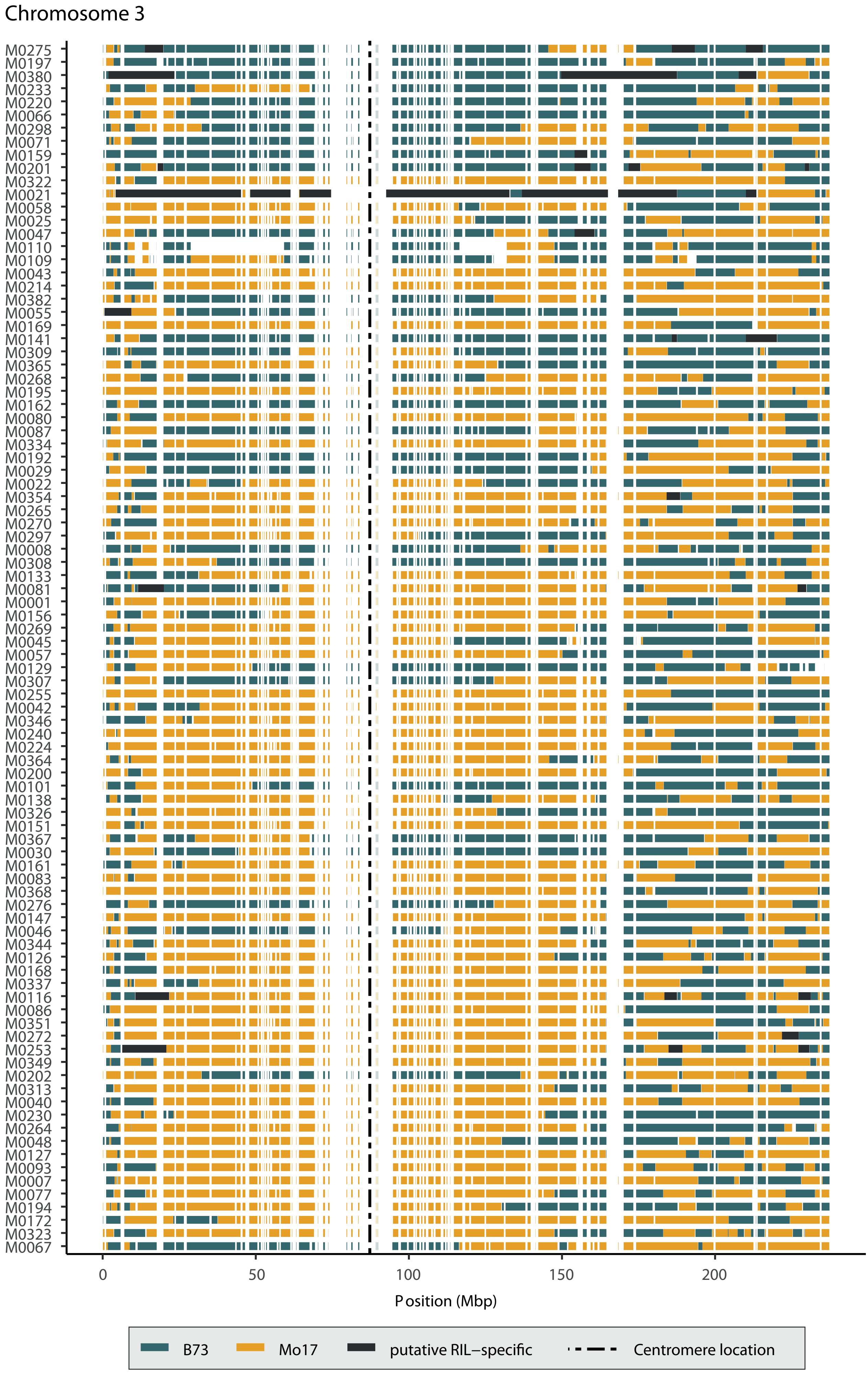

Chromosome 3
M0275
M0197
M0380
M0233
M0220
M0066
M0298
M0071
M0159
M0201
M0322
M0021
M0058
M0025
M0047
M0110
M0109
M0043
M0214
M0382
M0055
M0169
M0141
M0309
M0365
M0268
M0195
M0162
M0080
M0087
M0334
M0192
M0029
M0022
M0354
M0265
M0270
M0297
M0008
M0308
M0133
M0081
M0001
M0156
M0269
M0045
M0057
M0129
M0307
M0255
M0042
M0346
M0240
M0224
M0364
M0200
M0101
M0138
M0326
M0151
M0367
M0030
M0161
M0083
M0368
M0276
M0147
M0046
M0344
M0126
M0168
M0337
M0116
M0086
M0351
M0272
M0253
M0349
M0202
M0313
M0040
M0230
M0264
M0048
M0127
M0093
M0007
M0077
M0194
M0172
M0323
M0067
0
50
100
150
200
Position (Mbp)
B73
Mo17
putative RIL–specific
Centromere location

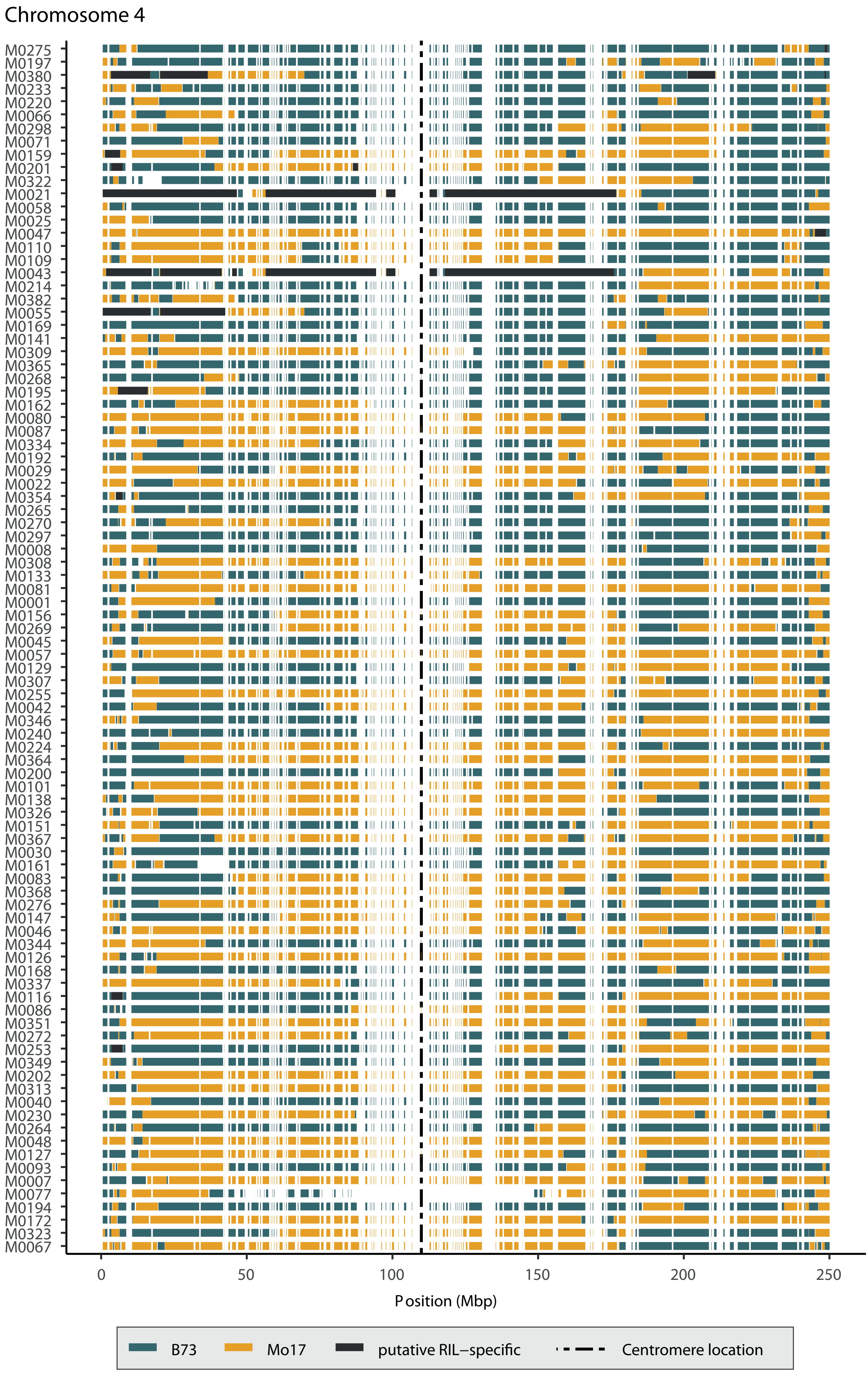
Chromosome 4
M0275
M0197
M0380
M0233
M0220
M0066
M0298
M0071
M0159
M0201
M0322
M0021
M0058
M0025
M0047
M0110
M0109
M0043
M0214
M0382
M0055
M0169
M0141
M0309
M0365
M0268
M0195
M0162
M0080
M0087
M0334
M0192
M0029
M0022
M0354
M0265
M0270
M0297
M0008
M0308
M0133
M0081
M0001
M0156
M0269
M0045
M0057
M0129
M0307
M0255
M0042
M0346
M0240
M0224
M0364
M0200
M0101
M0138
M0326
M0151
M0367
M0030
M0161
M0083
M0368
M0276
M0147
M0046
M0344
M0126
M0168
M0337
M0116
M0086
M0351
M0272
M0253
M0349
M0202
M0313
M0040
M0230
M0264
M0048
M0127
M0093
M0007
M0077
M0194
M0172
M0323
M0067
0
50
100
150
200
250
Position (Mbp)
B73
Mo17
putative RIL–specific
Centromere location

Chromosome 5

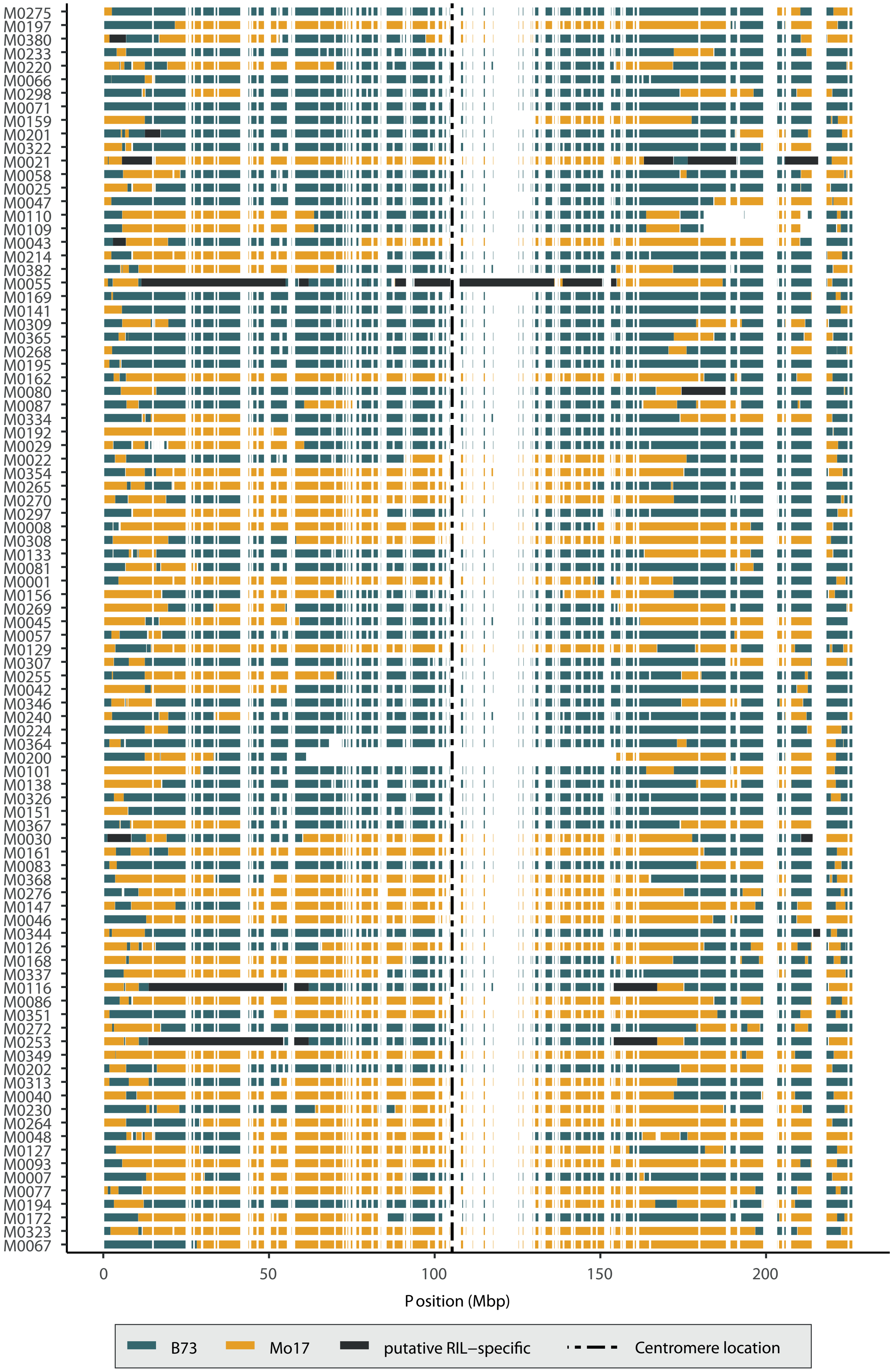

### Chromosome 6

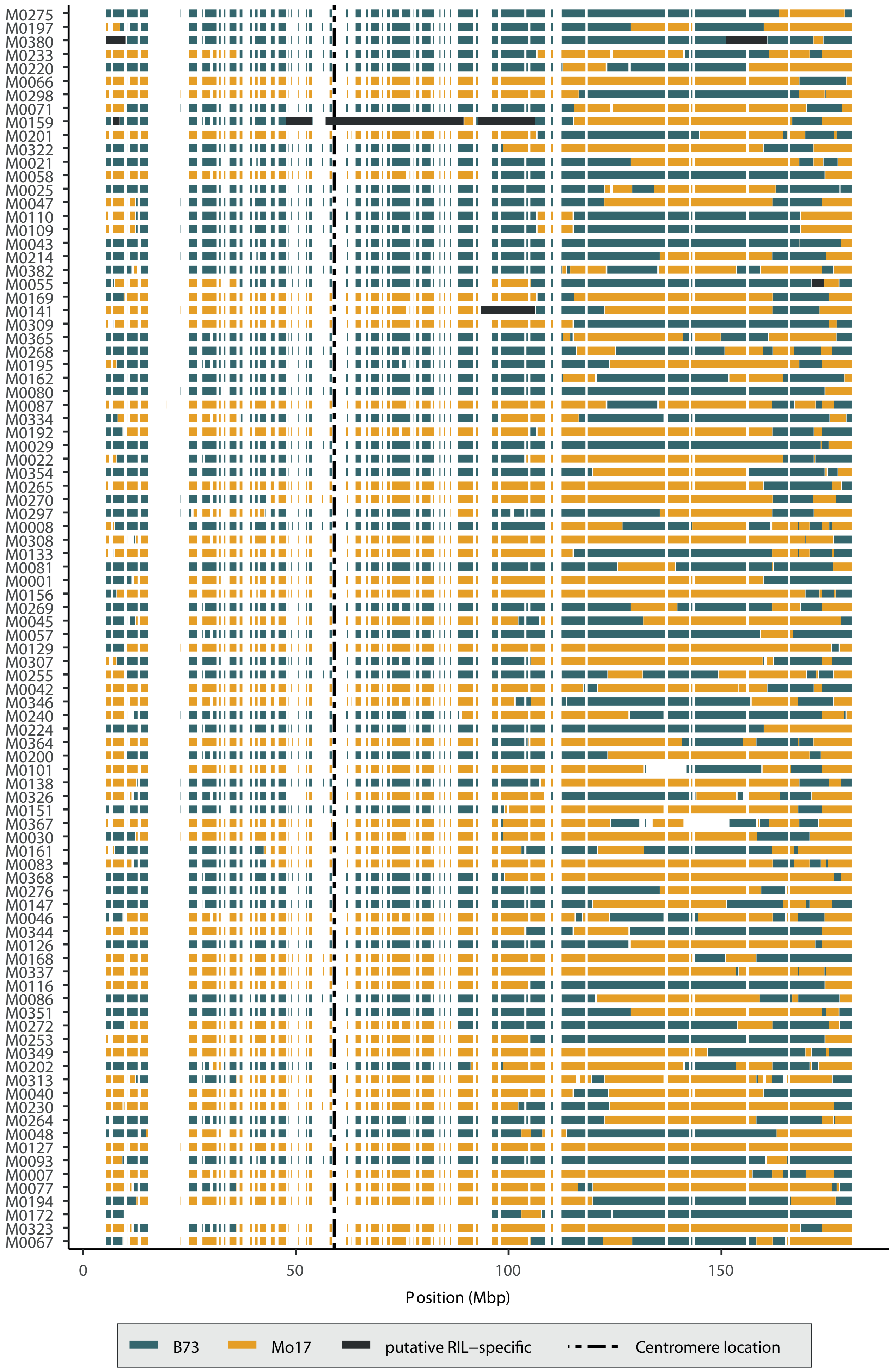

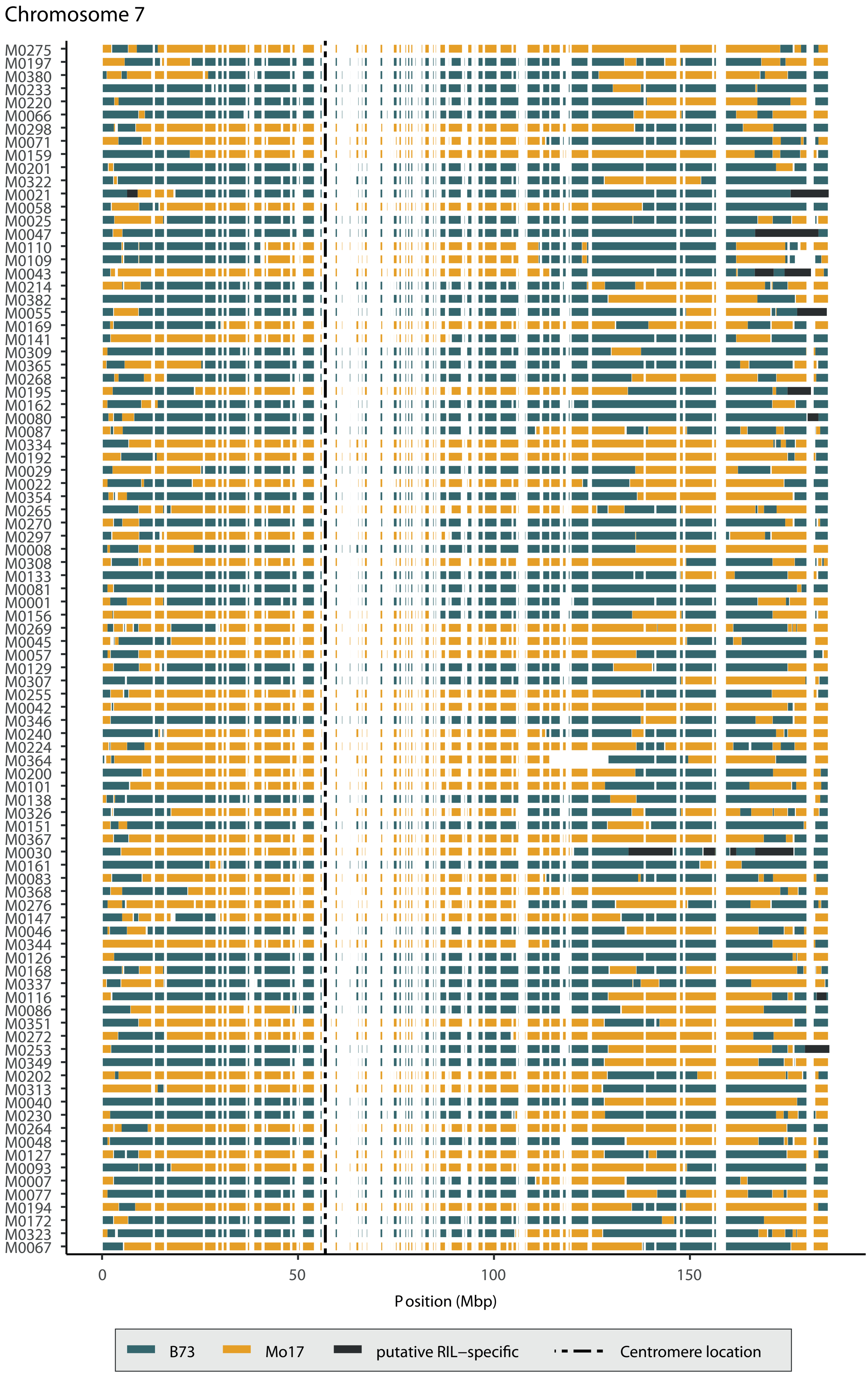
Chromosome 7
M0275
M0197
M0380
M0233
M0220
M0066
M0298
M0071
M0159
M0201
M0322
M0021
M0058
M0025
M0047
M0110
M0109
M0043
M0214
M0382
M0055
M0169
M0141
M0309
M0365
M0268
M0195
M0162
M0080
M0087
M0334
M0192
M0029
M0022
M0354
M0265
M0270
M0297
M0008
M0308
M0133
M0081
M0001
M0156
M0269
M0045
M0057
M0129
M0307
M0255
M0042
M0346
M0240
M0224
M0364
M0200
M0101
M0138
M0326
M0151
M0367
M0030
M0161
M0083
M0368
M0276
M0147
M0046
M0344
M0126
M0168
M0337
M0116
M0086
M0351
M0272
M0253
M0349
M0202
M0313
M0040
M0230
M0264
M0048
M0127
M0093
M0007
M0077
M0194
M0172
M0323
M0067
0
50
100
150
Position (Mbp)
B73
Mo17
putative RIL−specific
Centromere location

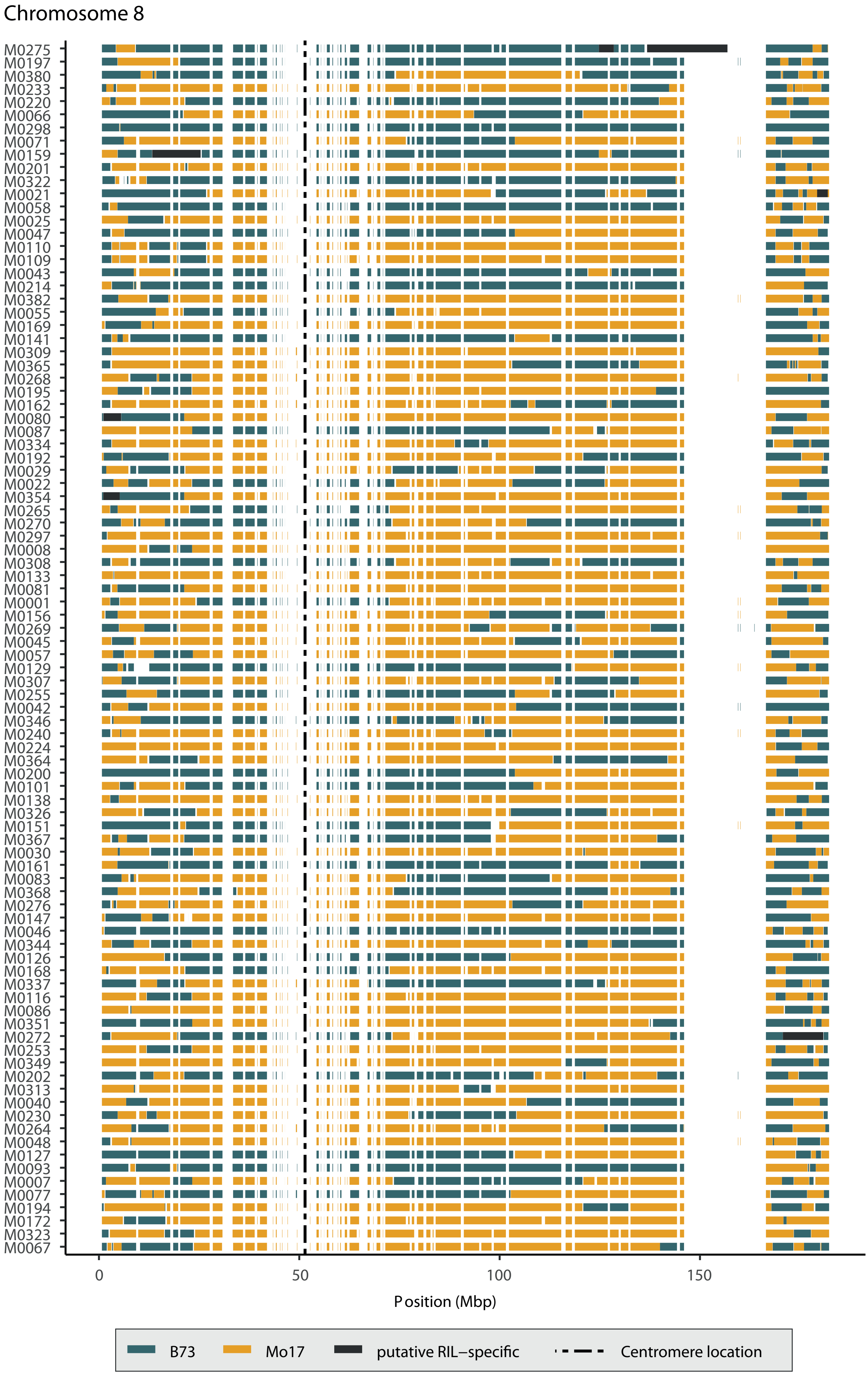
Chromosome 8
M0275
M0197
M0380
M0233
M0220
M0066
M0298
M0071
M0159
M0201
M0322
M0021
M0058
M0025
M0047
M0110
M0109
M0043
M0214
M0382
M0055
M0169
M0141
M0309
M0365
M0268
M0195
M0162
M0080
M0087
M0334
M0192
M0029
M0022
M0354
M0265
M0270
M0297
M0008
M0308
M0133
M0081
M0001
M0156
M0269
M0045
M0057
M0129
M0307
M0255
M0042
M0346
M0240
M0224
M0364
M0200
M0101
M0138
M0326
M0151
M0367
M0030
M0161
M0083
M0368
M0276
M0147
M0046
M0344
M0126
M0168
M0337
M0116
M0086
M0351
M0272
M0253
M0349
M0202
M0313
M0040
M0230
M0264
M0048
M0127
M0093
M0007
M0077
M0194
M0172
M0323
M0067
0
50
100
150
Position (Mbp)
B73
Mo17
putative RIL−specific
Centromere location

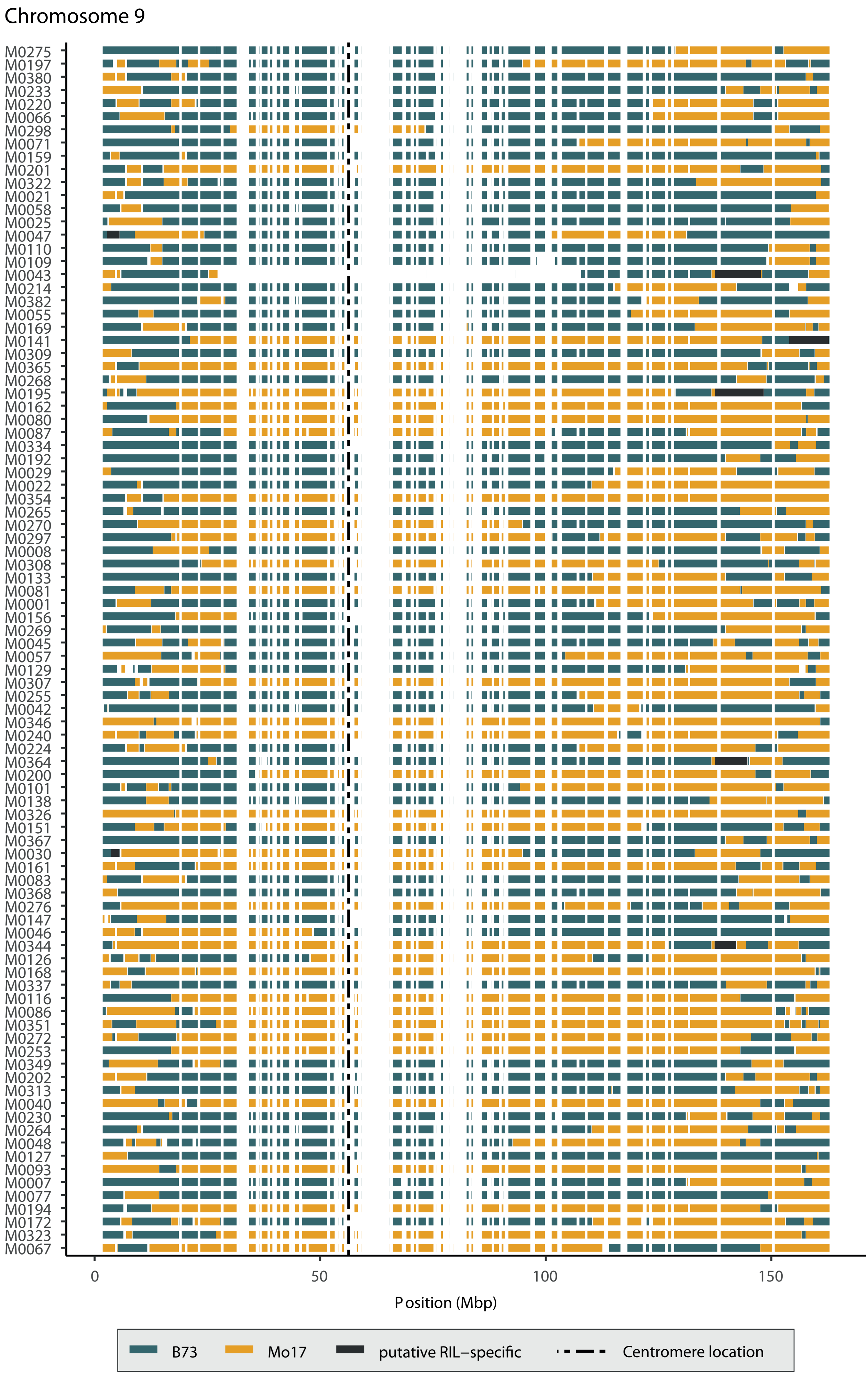

Chromosome 9
M0275
M0197
M0380
M0233
M0220
M0066
M0298
M0071
M0159
M0201
M0322
M0021
M0058
M0025
M0047
M0110
M0109
M0043
M0214
M0382
M0055
M0169
M0141
M0309
M0365
M0268
M0195
M0162
M0080
M0087
M0334
M0192
M0029
M0022
M0354
M0265
M0270
M0297
M0008
M0308
M0133
M0081
M0001
M0156
M0269
M0045
M0057
M0129
M0307
M0255
M0042
M0346
M0240
M0224
M0364
M0200
M0101
M0138
M0326
M0151
M0367
M0030
M0161
M0083
M0368
M0276
M0147
M0046
M0344
M0126
M0168
M0337
M0116
M0086
M0351
M0272
M0253
M0349
M0202
M0313
M0040
M0230
M0264
M0048
M0127
M0093
M0007
M0077
M0194
M0172
M0323
M0067
0
50
100
150
Position (Mbp)
B73
Mo17
putative RIL−specific
Centromere location

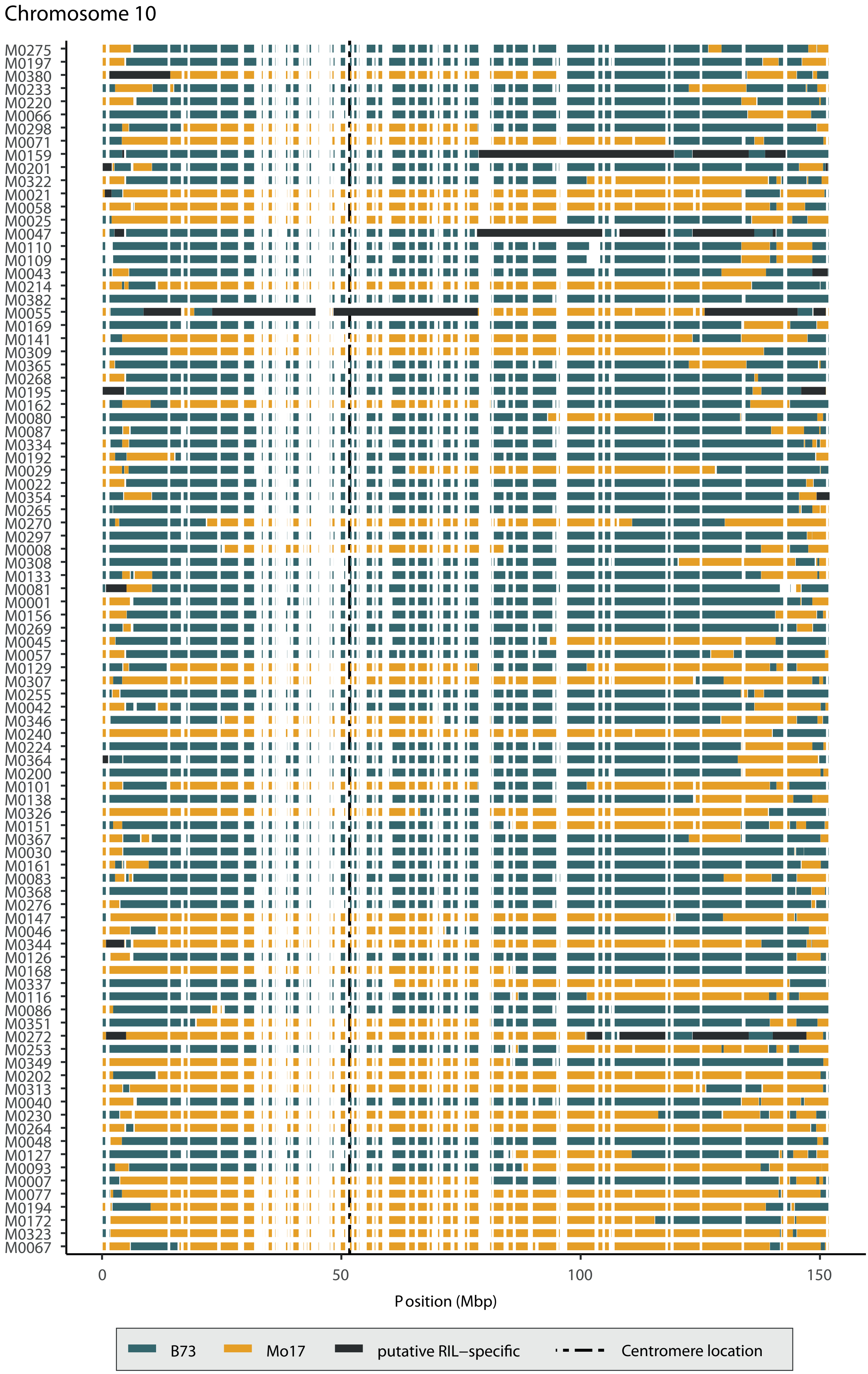

Chromosome 10
M0275
M0197
M0380
M0233
M0220
M0066
M0298
M0071
M0159
M0201
M0322
M0021
M0058
M0025
M0047
M0110
M0109
M0043
M0214
M0382
M0055
M0169
M0141
M0309
M0365
M0268
M0195
M0162
M0080
M0087
M0334
M0192
M0029
M0022
M0354
M0265
M0270
M0297
M0008
M0308
M0133
M0081
M0001
M0156
M0269
M0045
M0057
M0129
M0307
M0255
M0042
M0346
M0240
M0224
M0364
M0200
M0101
M0138
M0326
M0151
M0367
M0030
M0161
M0083
M0368
M0276
M0147
M0046
M0344
M0126
M0168
M0337
M0116
M0086
M0351
M0272
M0253
M0349
M0202
M0313
M0040
M0230
M0264
M0048
M0127
M0093
M0007
M0077
M0194
M0172
M0323
M0067
0
50
100
150
Position (Mbp)
B73
Mo17
putative RIL−specific
Centromere location

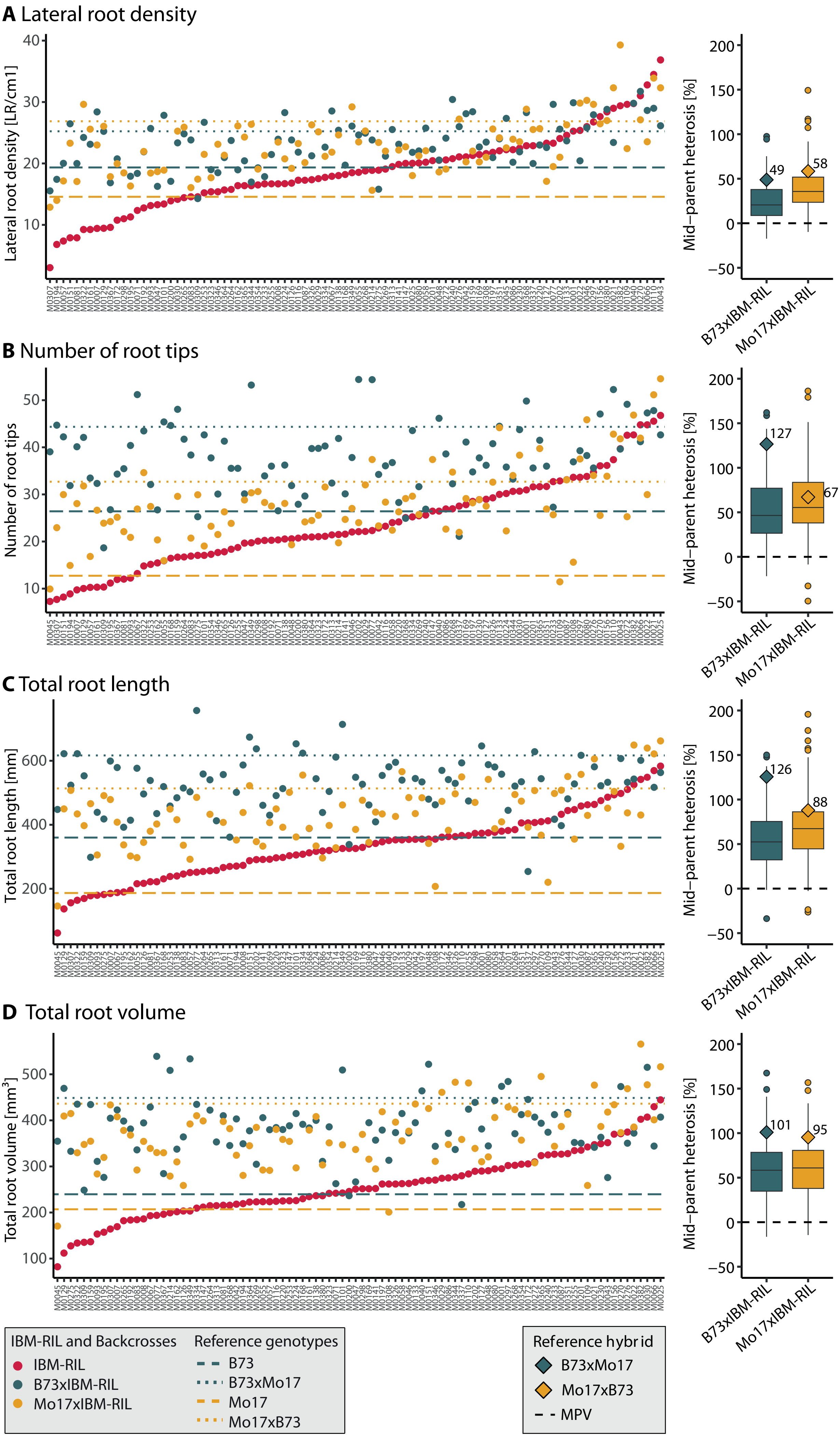
A Lateral root density
Lateral root density [LR/cm1]
B Number of root tips
Number of root tips
C Total root length
Total root length [mm]
D Total root volume
Total root volume [mm³]
Mid−parent heterosis [%]
B73xIBM-RIL
Mo17xIBM-RIL
49
58
127
67
126
88
101
95
IBM-RIL and Backcrosses
IBM-RIL
B73xIBM-RIL
Mo17xIBM-RIL
Reference genotypes
B73
B73xMo17
Mo17
Mo17xB73
Reference hybrid
B73xMo17
Mo17xB73
MPV

**Figure S3**: Phenotypic values and mid-parent heterosis (MPH) of **A** lateral root density, **B** Number of root tips, **C** Total root length, **D** Total root volume. On the left panel, the estimated means for each genotype are shown. The IBM-RILs (red) and the B73xIBM-RIL (blue) and Mo17xIBM-RIL (yellow) backcross hybrids are shown as points. The reference genotypes B73 (blue) and Mo17 (yellow) are shown as dashed lines and the reciprocal reference hybrids B73xMo17 (blue) and Mo17xB73 (yellow) as dotted lines. On the right panel, the mid-parent heterosis in percent of the parental mean is shown as boxplots for B73xIBM-RILs and Mo17xIBM-RILs. The MPH for the reference hybrids B73xMo17 (blue) and Mo17xB73 (yellow) is shown as diamond shaped points and their exact values are indicated. The dashed line indicates an MPV of 0.

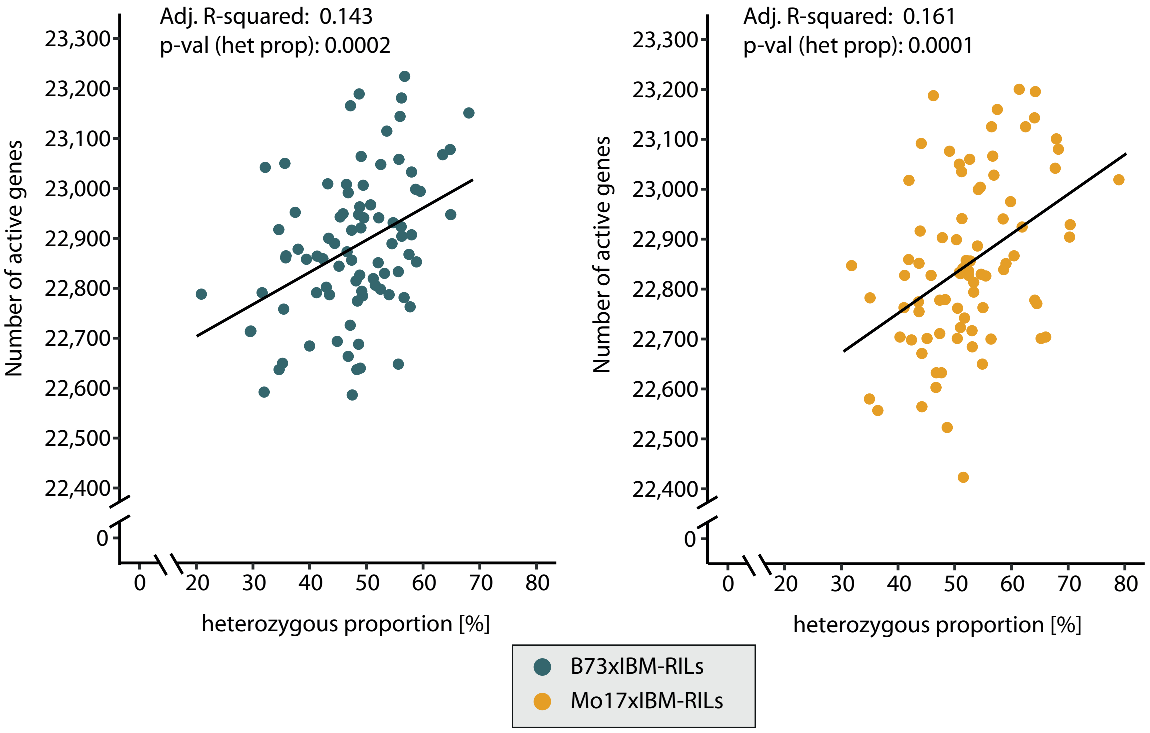


**Figure S4**: Correlation of heterozygosity and the total number of active genes in B73xIBM-RILs (left) and Mo17xIBM-RILs (right). The heterozygous proportion on the x-axis was calculated from the classified IBM-RIL regions. A linear regression with an intercept and the heterozygous proportion as covariate was fitted and the adjusted R-squared and p-value for the slope of the heterozygosity are indicated above the regression lines.

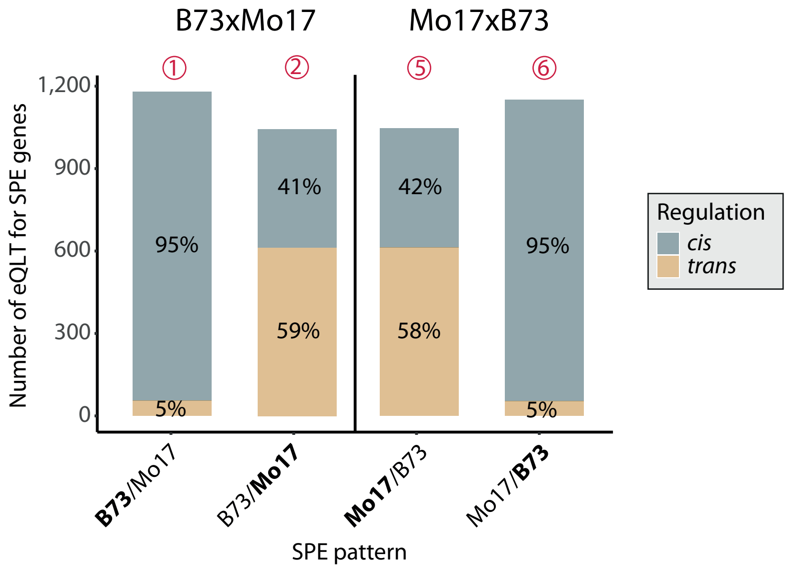


**Figure S5:** Regulation of SPE pattern genes in fully heterozygous reference hybrids B73xMo17 (left side) and Mo17xB73 (right side). The number of *cis*-(blue) and *trans*-(yellow) acting eQTL are given as bars, with the percentages of *cis* and *trans*-acting eQTL indicated per SPE pattern. The numbers above correspond to SPE pattern as indicated in Figure 2.

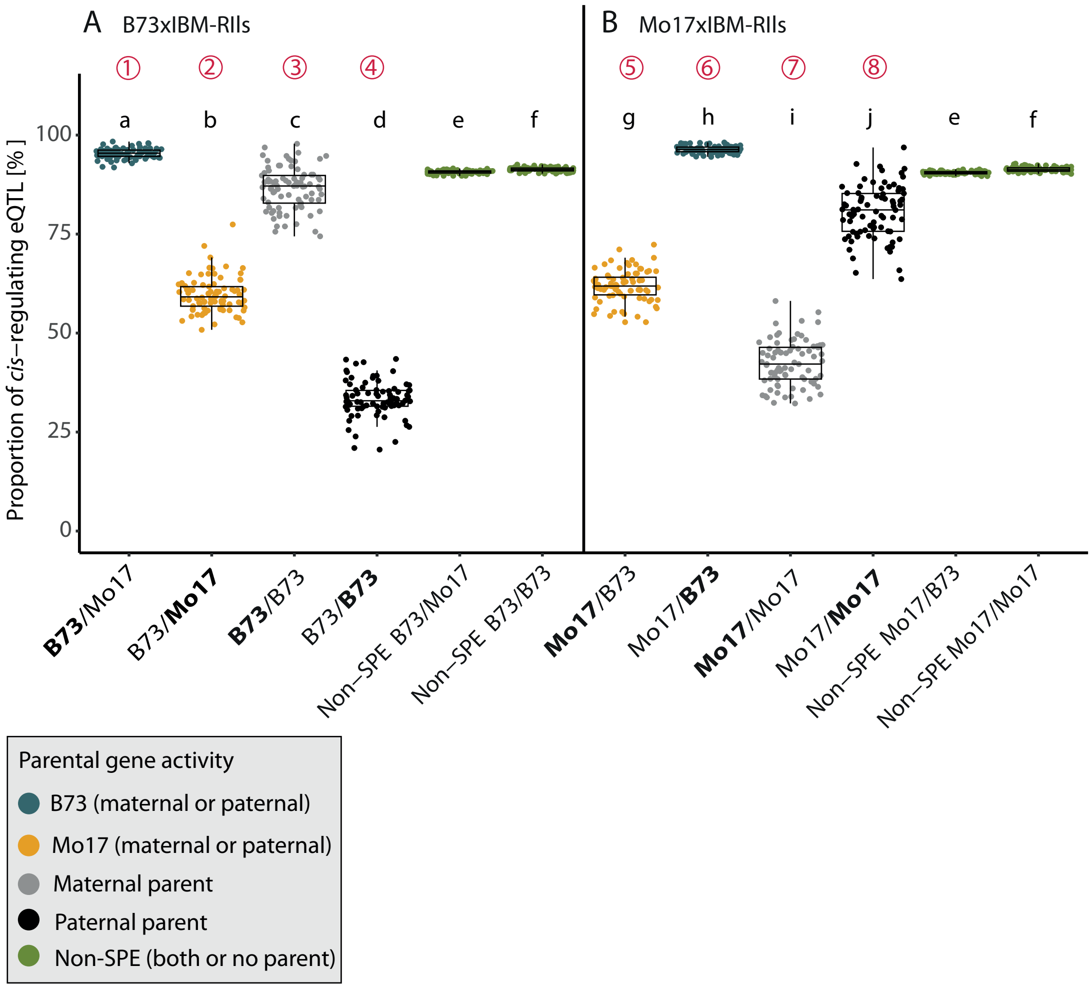


**Figure S6**: Proportion of *cis*- and *trans*- regulation in SPE pattern genes. Boxplots display the proportion of *cis*-regulation among SPE pattern and non-SPE pattern genes in the B73xIBM-RIL (**A**) and Mo17xIBM-RIL (**B**) hybrids. Different letters indicate significantly different proportions (a<0.05), identified with a gaussian mixed model with the hybrid as random effect, the SPE pattern and non-SPE pattern as a fixed factor and a diagonal variance component for the SPE and non-SPE pattern.

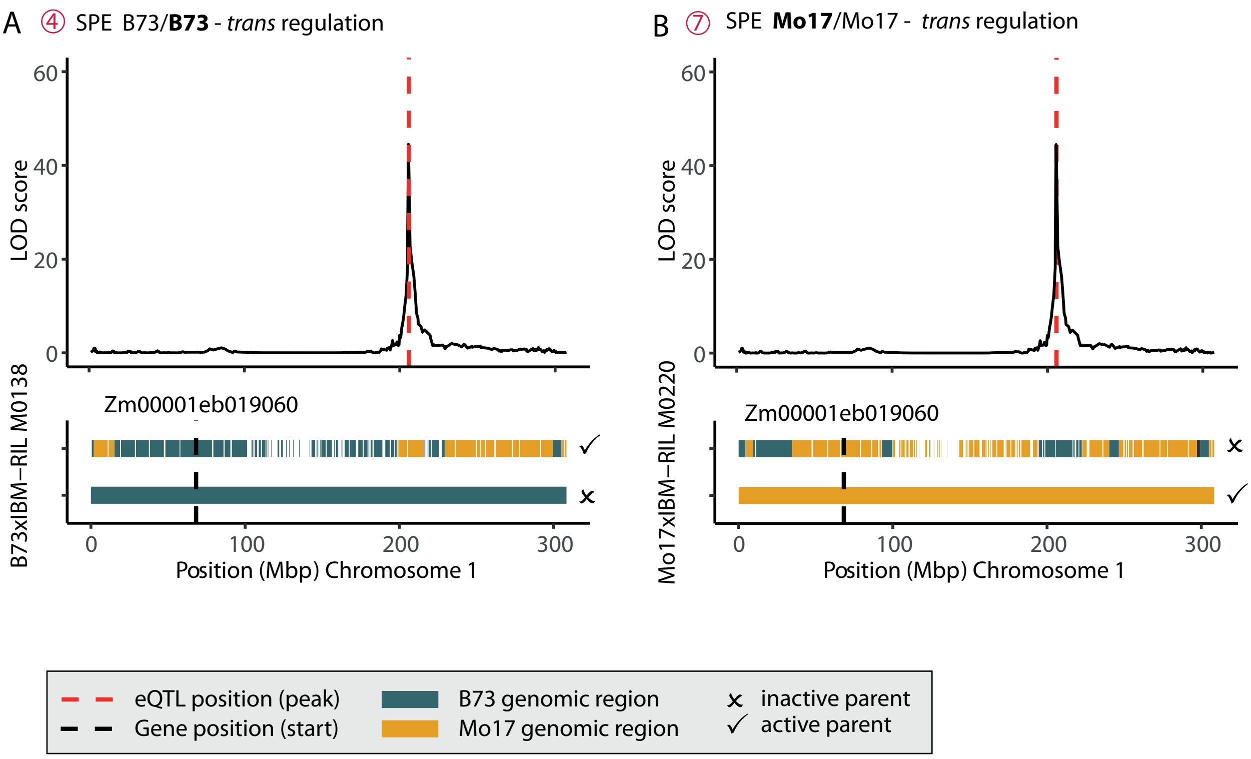


**Figure S7:** Examples for gene regulation pattern. **A** Homozygous SPE pattern 4 (B73/**B73**) gene in a B73xIBM-RIL, which is trans regulated from a heterozygous eQTL. **B** homozygous SPE pattern 7 (**Mo17**/Mo17) gene in a Mo17xIBM-RIL, which is trans-regulated homozygous gene from heterozygous eQTL.
