## Supplement Material and Methods SM1 and SM2 for "Regulation of heterosis-associated gene expression complementation in maize hybrids"

**Supplement Material SM1: Analysis of image-based phenotypic data and lateral root density**

After scanning the up to 8 maize seedlings per sample, the images were adjusted to have a minimum size of 600 x 600 px by adding a black frame around smaller images, using a custom Python script. The RootPainter software client (version 0.1.0) and server component (version 0.2.7) were used to train a convolutional neural network to recognize roots in images. The training dataset is a subset of the images, generated with the following settings: a maximum of 2 tiles per image, all images, and a target height and width of 700 pixels. The corrective annotation method of the software was used to mark the roots in the training dataset and the network was trained based on this data. Subsequently, all images were segmented using the trained model and converted to black recognized roots and white background[1]. The converted images were then analyzed in a batch using RhizoVision Explorer (version 2.0.3). The software was set to analyze only the largest root component to exclude roots from neighboring plants that could not be removed by image cropping. However, this approach resulted in incomplete analysis of some images with gaps in roots. To address this issue, an additional run was performed with settings to include all root objects bigger than 60 $mm^2$ [2]. The images with different results were examined and the correct results were saved for further analysis. Among the measured root parameters are the total root lengths and volume of all roots and of roots specific customary diameter ranges. One customary diameter range was set to 2.5 and above, so that only the seed falls into this category. The length and volume of roots from this category were subtracted from the total root length and volume. In addition, the number of total root tips in each image was measured. Technical outliers, like images with a brown background instead of a blue one and images where the roots were covered by name tags, were removed from the data.

For measuring the lateral root density, the uppermost cm of the primary root with emerged lateral roots was collected of each seedling and stored in 80% ethanol until counting. For each root piece, the number of lateral roots per cm was counted and used as lateral root density.

**Supplement Material SM2: SNP calling between B73 and Mo17 for sample evaluation**

In short, the GATK HaplotypeCaller was used to identify variants between the Mo17 samples and the B73v5 reference genome. Loci where the B73 samples also showed a variant were excluded. The resulting SNPs were compiled into a list of potential SNP loci. The aligned reads of each RNA-seq sample were then compared to this list, and the frequency of the B73 and Mo17 alleles at each SNP locus was determined ([3] adapted from[4]). The resulting allele frequencies were utilised to identify high quality samples.

In detail, to prepare the list of potential SNP loci, all Mo17 samples were combined into one file, using samtools merge and an index was created after renaming the RGSM (sample name field of readgroup information) for all samples to Mo17. The GATK HaplotypeCaller was run, setting the standard-min-confidence-threshold-for-calling to 20 and dont-use-soft-clipped-bases to true. This was done in parallel on each chromosome, by specifying intervals of whole chromosomes. The chromosome data was combined using GATK GatherVcfs, resulting in a file of Mo17 variants *vs* the B73v5 reference genome. The same was done for the B73 samples *vs* the B73v5 reference genome, because the B73 samples used in this study, might not be completely identical with the reference genome. These differences should be ignored. Therefore, the locations, where the B73 samples also show variance to the B73 genome were excluded from the list of possible SNP locations. This was done by a custom python script (Python3) [3]. The list of Mo17 variants was further filtered to contain only SNPs (no InDels) and only those SNPs which are homozygous for the SNP allele (putative Mo17) with bcftools view -i 'TYPE="snp" && GT="AA"' (from htslib version 1.14) [5]

To determine the allele frequencies of the B73 and Mo17 alleles at each of the SNP loci, all alignment files, (prepared as described in Material and Methods) were separately analyzed by a custom python script (Python2.7). The script checks at each identified Mo17 SNP position, how often the reference (putative B73) allele is present in the alignments and how often the SNP allele (putative Mo17) is present at the same site. These counts were then filtered for loci within protein-coding genes. The protein-coding genes were extracted from the maize B73 annotation file (http://ftp.ensemblgenomes.org/pub/plants/release-52/gff3/zea_mays/Zea_mays.Zm-B73-REFERENCE-NAM-5.0.52.gff3.gz) via a custom python script (Python3)[3]. It had to be ascertained for the SNP loci in protein-coding genes, that the reported reference and SNP allele counts correspond to the B73 and Mo17 allele of the germplasm used in this study (reference = B73, SNP = Mo17). This was done by retaining only loci, where the in Mo17 samples the SNP count is strongly predominant and in B73 samples the reference counts are strongly predominant. Therefore, the average of the Mo17 allele count and B73 allele count across the Mo17 samples was calculated. Only loci with minimum 95% Mo17 counts (and max 5% B73) on average were kept. The same was done for the B73 samples, where loci with at least 95% B73 counts were kept. The loci were further checked to have no third allele. Additionally, only those loci, where the average of the reference count in B73 samples is larger than 0.25, are kept. Considering the 48 B73 samples, that means a total count of 12 within all samples. This is done, to make sure that the reference (B73) allele is confirmed by sufficient coverage in our samples. (The SNP allele is already confirmed, because if there was no coverage for it, the SNP could not have been called).

Due to some samples clustering in a different group (e.g. B73xIBM-RIL sample in Mo17xIBM-RIL group), visible in the PCA plot (Figure 1C), all samples were investigated to check, if their information is consistent within the three replicates of a genotype and a triplet. First, the homozygosity in the inbred IBM-RIL samples was calculated. Only loci with a minimum coverage of 0.5 counts per million (cpm) were considered. The percentage of homozygous B73 and Mo17 loci was calculated, whereas a locus was considered homozygous if 95% of all counts belong to the major allele. The sum of the percentages of homozygous B73 and Mo17 loci is calculated and is expected to be close to 100%. This was done to check whether all the pooled roots within an RNA-seq sample came from the correct genotype. Thus, samples with less than 95% homozygous loci were excluded. Furthermore, IBM-RILs with only one remaining replicate were excluded.

The remaining IBM-RIL samples were then compared to their B73xIBM-RIL and Mo17xRIL hybrid samples. A B73xIBM-RIL hybrid for example, should be homozygous at the loci where the IBM-RIL has the homozygous B73 allele and respectively, Mo17xIBM-RIL hybrids should be homozygous for the Mo17 allele at the loci where the IBM-RIL is homozygous Mo17. The loci were filtered for those with consistent allele information across the samples of the same IBM-RIL. Loci that were homozygous for Mo17 were selected in the Mo17xIBM-RIL hybrid of the same triplet and the homozygosity was calculated as described for the IBM-RIL samples. The same was done with B73 loci in the B73xIBM-RIL hybrids. Again, samples with less than 95% homozygous loci were excluded. In total, 175 samples were excluded, because they were not homozygous in at least 95% of supposedly homozygous loci. Beforehand, 10 samples had been excluded due to their library size of <5 million read counts. As a result, 17 samples were excluded because they were the only replicate left of the respective genotype. In our analyses, we always compared both parents and the resulting hybrid, thus 90 hybrid samples were excluded, because the respective paternal IBM-RIL samples were all excluded and 8 IBM-RIL samples were excluded, because all corresponding hybrid samples were excluded. A total of 852 samples remained for the final SNP calling of all samples against the B73v5 reference genome.
